# Evolution of an unstable sex determination system in white Guinea yam (*Dioscorea rotundata*)

**DOI:** 10.64898/2026.09.08.750133

**Authors:** Aoi Kudoh, Yu Sugihara, Kohtaro Iseki, Ryo Matsumoto, Koki Minoji, Kiyoshi Onai, Akira Abe, Satoshi Natsume, Toshiyuki Sakai, Kaori Oikawa, Motoki Shimizu, Kazue Itoh, Hiroaki Adachi, Kodai Honda, Shinsuke Yamanaka, Paterne Agre, Patrick Adebola, Asrat Asfaw, Ryohei Terauchi

## Abstract

Plants often produce unstable sex phenotypes; however, the underlying evolutionary and molecular mechanisms are unclear. In white Guinea yam (*Dioscorea rotundata*), the most economically important tuber crop, genetic improvement is critical for food security but remains constrained by dioecy and complex sex phenotypes. White Guinea yam has a ZZ/ZW sex determination system in which ZZ individuals are consistently male, whereas ZW individuals can be female, male, or monoecious. Here we show that chromosome 11 is the sex chromosome, featuring a 1.2-Mb inversion between the Z and W chromosomes. A 170-kb W-specific region carries the microRNA gene *dro-MIR432*, likely involved in sex switching. The recent emergence of this miRNA-mediated ZZ/ZW system, distinct from the XY/XX system within *Dioscorea*, underscores the dynamic evolution of plant sex determination.

---

Sexual reproduction is nearly universal among eukaryotes, and male and female sexual phenotypes are broadly observed across taxa (*1*). Flowering plants display diverse sexual reproduction strategies, including dioecy, in which male and female flowers are borne on separate individuals, and monoecy, in which male and female flowers are borne on the same individual (*2*). The molecular basis of genetically determined sex in dioecious and monoecious plants has begun to be elucidated (*3–5*). However, sex expression is not always stable and often changes within a single generation in diverse plant lineages. The evolution and molecular basis of such unstable sex expression are poorly understood (*6*).

The monocot genus *Dioscorea*, in the family Dioscoreaceae, contains approximately 630 species, including economically important yam species that are cultivated worldwide (*7*). Global yam production reached ∼92.4 million tons in 2024; however, productivity per unit area has stagnated (*8*), raising concerns about meeting future demand in yam-consuming regions. Therefore, genetic improvement of yam is critical to ensure food security (*9*). A major constraint on yam breeding is its dioecy, which prevents the fixation of desirable traits through self-pollination (*10*). Recent genomic studies have begun to reveal the molecular basis of genetic sex determination in *Dioscorea*. The wild yam *D. tokoro*, which is endemic to East Asia, has a male heterogametic sex-determination system (XY/XX). Two Y-specific genes are likely involved in sex determination: *BLH9*, encoding a homeobox protein, and *HSP90*, encoding a molecular chaperone (*11*). Water yam (*D. alata*), which is widely cultivated in pantropical regions, also has an XY/XX system; its putative sex-determining region was localized to a ∼7.6-Mb region of the Y chromosome (*12*).

Here, we explored the molecular basis of sex determination in white Guinea yam (*D. rotundata*), which is mainly cultivated in West and Central Africa, a region accounting for over 90% of worldwide yam production (*8*, *10*, *13*, *14*). A previous linkage analysis revealed that white Guinea yam has a female heterogametic sex-determination (ZZ/ZW) system and identified the female-specific DNA marker *sp16* in the putative W-linked genomic region (*15*). Genotyping using the *sp16* marker revealed that individuals with the ZZ genotype are consistently male, whereas individuals with the ZW genotype display unstable sex phenotypes, including the occasional production of male flowers on otherwise female individuals (*15*) (Fig. 1A–C). The variability of sex phenotypes was also reported in four-year field trials, where ZW individuals exhibited male, female, and monoecious phenotypes, with the same individual showing different sex phenotypes across years (*16*) (Fig. 1D). To elucidate the sex determination system in white Guinea yam, we generated a haplotype-resolved genome assembly of this species. Together with transcriptome analysis, this assembly identified a W-specific microRNA (miRNA) as a possible sex determinant likely involved in the unstable sex expression of this important crop.

**Fig. 1.**
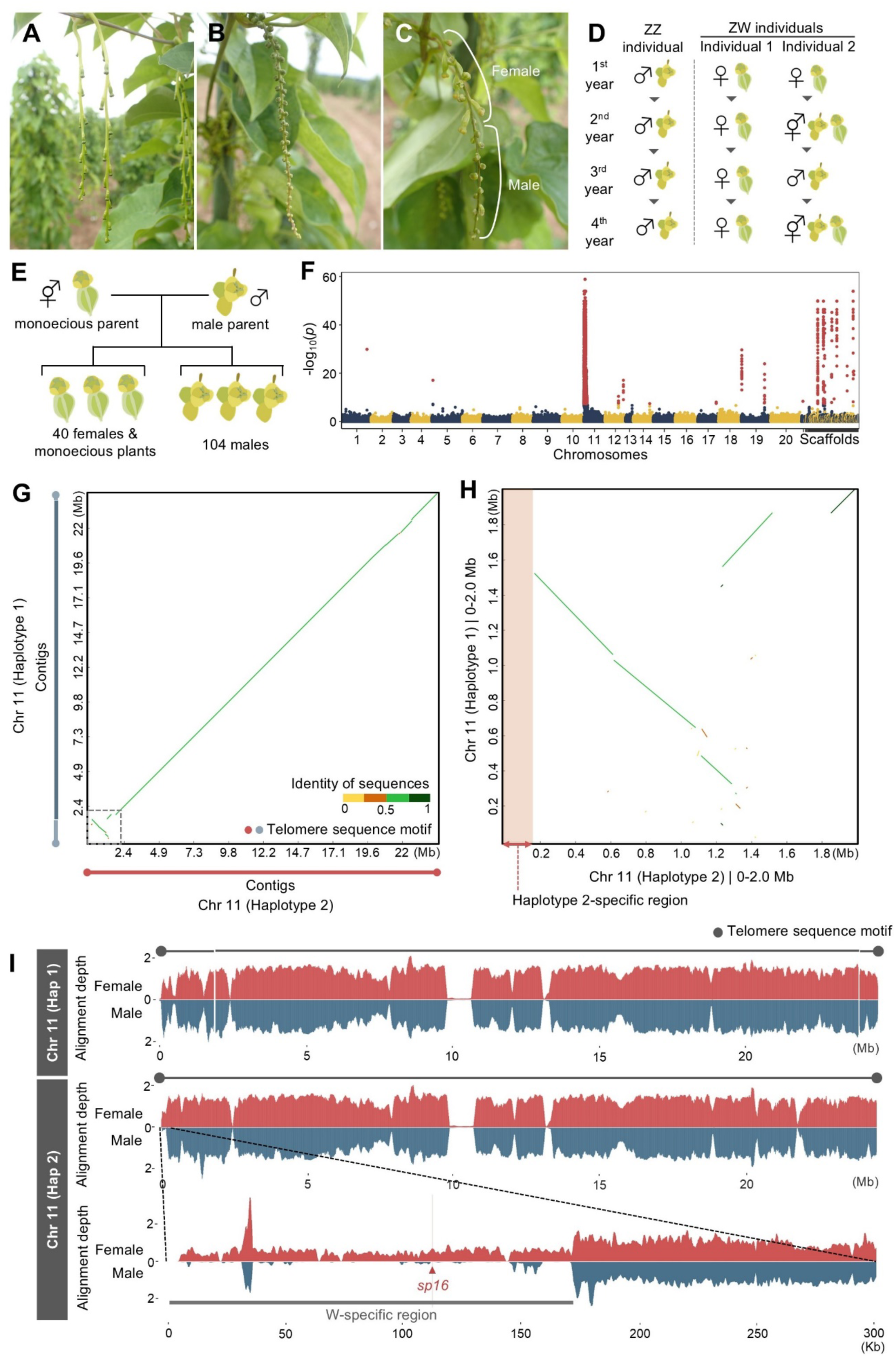
Haplotype-resolved genome assemblies, association analysis, and mapping coverage analysis reveal a W-specific region in the white Guinea yam *Dioscorea rotundata* genome. **(A–C)** Representative *D. rotundata* female **(A),** male **(B)**, and monoecious **(C)** inflorescences. **(D)** Diagram of variability in sex phenotype, adapted from Iseki et al. (2022). Individuals with the ZZ genotype show consistent maleness, whereas individuals with the ZW genotype exhibit male, female, and monoecious phenotypes, with sex phenotype varying across years. **(E)** F1 progeny, comprising 40 females and monoecious individuals and 104 males, used for association analysis to identify genomic regions associated with the sex phenotype. **(F)** Manhattan plot of the genome-wide association study (GWAS) results based on the genotypes and sex phenotypes of F1 individuals. Red points indicate single nucleotide polymorphism (SNP) markers significantly associated with the sex phenotype. A significance threshold of 5% false discovery rate adjusted by Bonferroni correction was applied. **(G)** Alignment-based dot plot comparing the assemblies of chromosome 11 from haplotype 1 and chromosome 11 from haplotype 2. Blue lines on the left and red lines at the bottom indicate the positions of the contigs for haplotype 1 and haplotype 2, respectively. The dotted square indicates the region shown in detail in (H). **(H)** Detailed view of the 2-Mb distal end of the Z and W chromosomes. The double-sided arrow indicates a 170-kb haplotype 2-specific region. **(I)** Alignment depth analysis for chromosome 11 from the haplotype 1 and 2 assemblies. Female and male mapping depths are shown in red and blue, respectively, based on Illumina short reads from the female individual TDr97_00917 and the male individual TDr97_00777 mapped to the haplotype 1 and 2 assemblies. The top plot shows chromosome 11 from haplotype 1, defined as the putative Z chromosome; the middle plot shows chromosome 11 from haplotype 2, defined as the putative W chromosome; and the bottom plot shows a detailed view of the indicated region in the middle plot, including the putative W-specific region and the female-specific DNA marker *sp16*.

## A W-specific region

Building on the putative W-linked region on chromosome 11 previously associated with sex by bulked segregant analysis (BSA) (*15*), we performed a genome-wide association study (GWAS) to further delineate the sex-associated region. We analyzed 156 F1 progeny derived from a cross between a monoecious parent and a male parent, comprising 40 females or monoecious plants, 104 males, and 12 non-flowering plants (Fig. 1E). Using Illumina short-read sequences from the F1 progeny aligned to the published consensus genome assembly (*13*), we examined the association between single-nucleotide polymorphism (SNP) genotypes and sex phenotypes. The GWAS revealed significant associations between genotypes at the distal end of chromosome 11 and sex phenotypes, consistent with the W-linked region previously identified by BSA (*15*) (Fig. 1F).

To characterize the genomic structure of the sex-associated region at the distal end of chromosome 11 in more detail, we reconstructed haplotype-resolved genome assemblies from the monoecious parent TDr96_F1 (Text S1; fig. S1; Table S1). Alignment-based dot plot comparison of the haplotype 1 and haplotype 2 assemblies identified an ∼1.2-Mb inverted region at the distal end of chromosome 11 (Fig. 1G), adjacent to a 170-kb haplotype 2–specific region (Fig. 1H).

To characterize this haplotype 2–specific region, we mapped Illumina short-read sequences from female and male individuals to the new assemblies and evaluated normalized read depth across the region. This 170-kb region was covered at a depth of 0.5× in female sequences but showed almost no (0×) depth in male sequences (Fig. 1I). The region contains the *sp16* marker previously identified within the putative W-linked region (*15*).

Based on these results, we defined chromosome 11 from haplotype 1 as the Z chromosome and chromosome 11 from haplotype 2 as the W chromosome. The Z and W chromosomes are differentiated by a ∼170-kb W-specific region and an adjacent ∼1.2-Mb telomeric inversion. This inversion may act as an effective barrier to recombination between the Z and W chromosomes. The W-specific region contains 17 predicted protein-coding genes (Table S2) and a candidate microRNA gene (Table S3). A sequence similarity search of the *D. rotundata* genome using the 17 predicted protein-coding genes identified low-identity homologs of these genes distributed across multiple autosomes of the white Guinea yam genome, rather than clustered in a specific genomic region. This observation suggests that the W-specific region may have formed through gene loss from the Z chromosome or gene accumulation from multiple autosomes, rather than through a recent duplication of a genomic region (Table S4).

## A W-specific miRNA

The presence of a W-specific region, together with the observation that ZW individuals can bear both female and male flowers, suggests that the W-specific region may carry a sex-determining gene whose activity varies, resulting in unstable sex expression. To identify candidate genes involved in sex determination (Fig. 2A), we conducted transcriptome analyses using RNA-seq and small RNA-seq data from female and male flowers at three developmental stages (Fig. 2B).

**Fig. 2.**
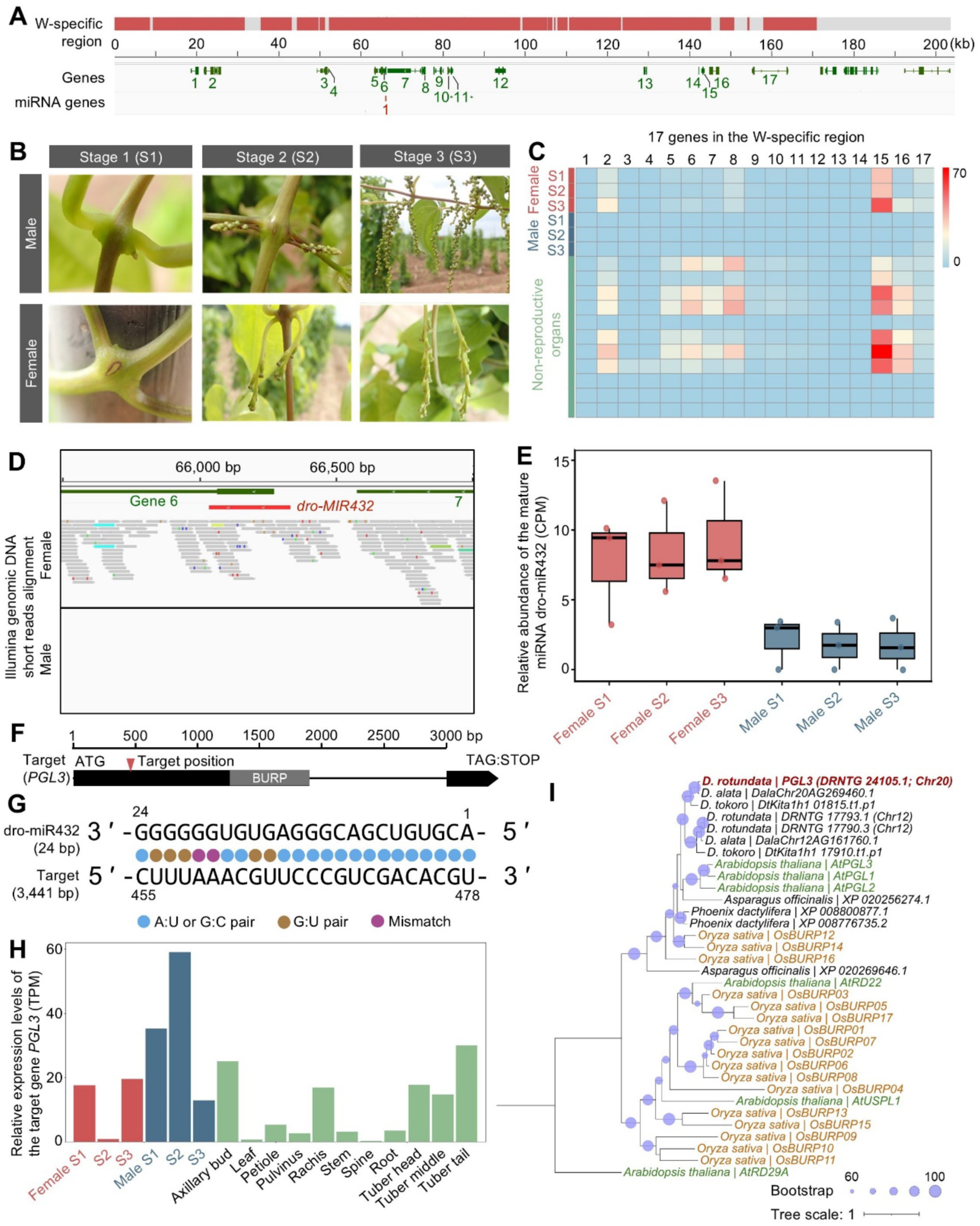
The W-specific miRNA dro-miR432 is produced specifically during the early stage of female flower development and potentially targets *PGL3*. **(A)** The W-specific region contains 17 genes and one locus for miRNA gene. **(B)** Representative photographs showing the three stages of male and female flower development used in transcriptome analysis. **(C)** Heatmap representation of the expression levels of 17 genes in the W-specific region. Expression levels are shown as normalized transcripts per million (TPM) values. **(D)** Coverage of Illumina short-read genomic sequencing data from male and female individuals across the *dro-MIR432* locus. **(E)** Relative levels of the mature miRNA dro-miR432 during the three stages of male and female flower development. The box boundaries represent the first (Q1) and third (Q3) quartiles, the horizontal line within each box indicates the median, and the whiskers extend to the most extreme data points within 1.5 × the interquartile range (IQR). **(F)** Gene structure of the target gene *PGL3*. Black boxes indicate coding regions; the sequence encoding the BURP domain is shown in gray. The red triangle indicates the predicted target site of dro-miR432. **(G)** Sequence alignment between dro-miR432 and its predicted target site in *PGL3*. **(H)** Relative *PGL3* expression levels during the three stages of male and female flower development and in non-reproductive organs. Each developmental stage is represented by a single biological sample. **(I)** Phylogenetic tree of PGL3 from white Guinea yam and BURP domain–containing proteins from *Oryza sativa*, *Arabidopsis thaliana*, and various monocotyledonous species (*Asparagus officinalis*, *Dioscorea tokoro*, *Dioscorea alata*, and *Phoenix dactylifera*). The circles in the tree nodes indicate bootstrap values. Tree scale indicates amino-acid substitutions per site.

The W-specific region contained 17 predicted protein-coding genes (Fig. 2A), including seven genes encoding proteins with functional annotations based on significant BLASTX hits to proteins in the Swiss-Prot database, and ten genes encoding proteins without functional annotations (Table S2). None of these seven proteins has previously been implicated in floral development. Few, if any, RNA-seq reads from male flowers were mapped to the 17 genes, as expected (Fig. 2C), and no gene showed significantly higher expression in female flowers compared to non-reproductive organs (Table S2).

Most miRNA genes are transcribed into a long transcript called the primary microRNA transcript (pri-miRNA), which is processed to generate functionally mature miRNAs typically ∼22 nucleotides long (*17*, *18*). The W-specific region contains a female-specific candidate miRNA gene, as supported by genomic read coverage in a female individual but not in a male individual (Fig. 2D). This gene is predicted to be transcribed into the pri-miRNA pri-miR432 (Fig. 2A). The 24-nt mature miRNA dro-miR432, predicted to be processed from pri-miR432, was significantly more abundant in female flowers than in male flowers at all three developmental stages (Fig. 2E; Table S3). miRNAs function by binding to the mRNAs of target protein-coding genes and repressing their expression via mRNA cleavage and/or translational inhibition (*17*). dro-miR432 is predicted to target transcripts of *PGL3*, a gene located on chromosome 20 (Fig. 2F; fig. S2). Sequence alignment between dro-miR432 and its predicted target site in *PGL3* revealed 17 A:U or G:C base pairs and five G:U wobble pairs (Fig. 2G; fig. S2). The largest difference in *PGL3* expression between female and male flowers was observed at stage 2, when expression was markedly lower in female flowers but reached its highest level in male flowers (Fig. 2H).

*PGL3* is closely related to the *Arabidopsis thaliana* genes *POLYGALACTURONASE 1 BETA-LIKE PROTEIN 1* (*AtPGL1*), *AtPGL2,* and *AtPGL3*, which encode BURP domain–containing proteins, and the rice (*Oryza sativa*) genes *BURP DOMAIN-CONTAINING PROTEIN 12* (*OsBURP12*), *OsBURP14* and *OsBURP16* (Fig. 2I; Table S5). OsBURP12 has been reported to function in germinating rice seeds and seedlings, particularly in salinity-stress tolerance (*19*) and cold-stress responses (*20*). In both studies, *OsBURP12* knockout lines showed significantly lower polygalacturonase activity (*19*, *20*). Polygalacturonases are cell wall hydrolytic enzymes that participate in pollen development in diverse plant species. For example, multiple polygalacturonase genes known as *Brassica campestris Male Fertility* genes were identified in male-sterile mutants of Chinese cabbage (*21–24*). The polygalacturonase homolog *NO SPINE POLLEN* (*GhNSP*) functions in pollen exine formation in cotton, and the *Ghnsp* mutant exhibits male sterility (*25*). In the dioecious willow *Salix gilgiana*, polygalacturonases appear to be involved in pollen grain development during male gametogenesis (*26*). Therefore, *D. rotundata PGL3*, which shares sequence similarity with *OsBURP12,* might be required for successful male flower development in white Guinea yam through its association with polygalacturonase activity. These results suggest that the sex phenotype of white Guinea yam might be controlled by the presence or absence of the miRNA dro-miR432, leading to low or high expression of its target gene *PGL3*, respectively (Fig. 2H).

To validate the functional relevance of dro-miR432 and its target gene *PGL3,* we performed a dual-luciferase assay, a technique commonly used for functional validation of miRNAs in animals (*27*, *28*) and plants (*29*, *30*). Briefly, we designed a dual-luciferase reporter plasmid containing a *Renilla* luciferase (R-Luc) expression cassette, which functions as an internal control to standardize expression, and a firefly luciferase (F-Luc) expression cassette containing the predicted target sequence between F-Luc and the 3’ untranslated region (Fig. 3A). We co-infiltrated *Nicotiana benthamiana* leaves with this dual-luciferase reporter construct and a plasmid driving expression of the pri-miRNA (Fig. 3B). To evaluate targeting of the reporter by the candidate miRNA, we used three plasmids for co-infiltration: a control harboring the β-glucuronidase (*GUS*) gene, the native pri-miR432 sequence (miRNA; Fig. 3C), and a mutated pri*-*miR432 sequence (Mutated; Fig. 3D). The mutated pri-miR432 sequence retained the nucleotide composition of the native miRNA-targeting sequence while randomizing nucleotide order.

**Fig. 3.**
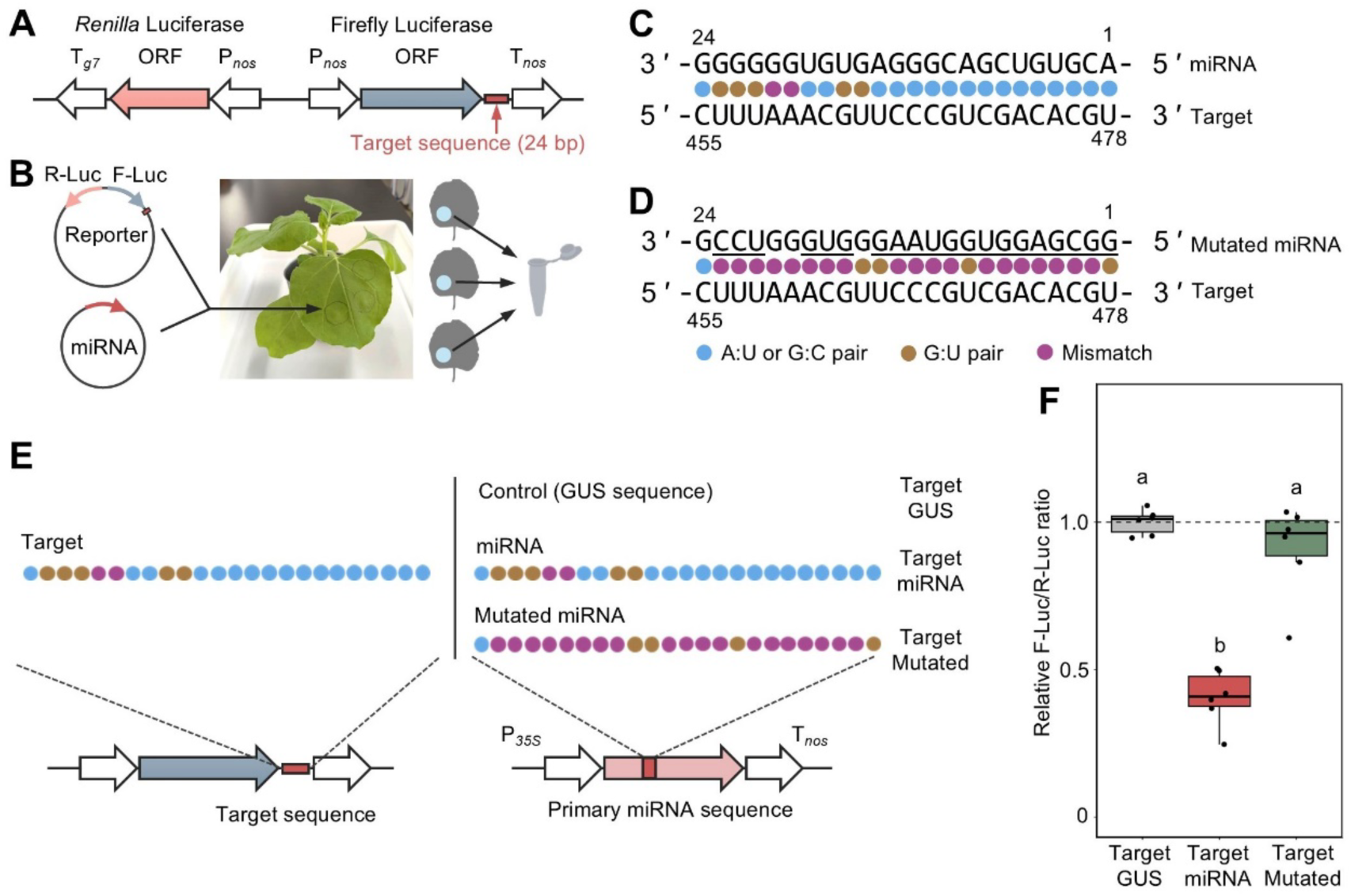
The candidate miRNA dro-miR432 targets its predicted binding sequence in *PGL3* in a dual-luciferase reporter assay in *Nicotiana benthamiana* leaves. **(A)** Design of the plasmid for the dual-luciferase reporter containing two luciferase expression cassettes: a *Renilla reniformis* luciferase expression cassette (internal control to standardize expression) and a firefly luciferase expression cassette (used to test the putative miRNA target sequence from *PGL3*). **(B)** Diagram of the Agrobacterium tumefaciens–mediated infiltration assay in *N. benthamiana* leaves. A mixture of *Agrobacterium* cultures carrying a dual-luciferase reporter plasmid or a pri-miRNA expression plasmid was co-infiltrated into *N. benthamiana*. Following incubation, the leaf tissues were collected and used for the dual-luciferase reporter assay. **(C)** The native core sequence for recognizing targets in the primary miRNA pri*-*miR432 (miRNA) and the putative miRNA target sequence from *PGL3* in the reporter plasmid (Target). **(D)** The mutated core sequence in pri*-*miR432mut (Mutated miRNA) and the putative miRNA target sequence from *PGL3* in the reporter plasmid (Target). The mutated core sequence contains the original nucleotide composition, but the sequence is randomized. **(E)** The reporter containing the target sequence from *PGL3* (left) was co-infiltrated with each of three expression constructs (right): (1) reporter and *GUS* expression construct (Target-GUS) as a control, (2) reporter and pri*-*miR432 expression construct (Target-miRNA), (3) reporter and mutated pri*-*miR432 expression construct (Target-Mutated) constructs. **(F)** Relative firefly luciferase to *Renilla* luciferase ratios (F-Luc/R-Luc) after 72 h ofincubation in the three sets of co-infiltrated leaves. The relative F-Luc/R-Luc ratios were compared using pairwise Welch’s *t*-tests with Holm-adjusted *p*-values. Three independent experiments were performed, and the results of one experiment are shown. The results of all three experiments are shown in fig. S3. The box boundaries represent the first (Q1) and third (Q3) quartiles, the horizontal line within each box indicates the median, and the whiskers extend to the most extreme data points within 1.5 × the interquartile range (IQR).

To assess the functional relevance of the miRNA target site, we performed three co-infiltration treatments in *N. benthamiana* leaves using the dual-luciferase reporter (Target) and one of three expression constructs. The three treatments were designated Target-GUS, Target-miRNA, and Target-Mutated (Fig. 3E). In the co-infiltration assays, the intact candidate miRNA significantly repressed relative F-Luc activity, whereas no repression was observed in the GUS control or mutated-miRNA treatment (Fig. 3F). Similar results were obtained in nine independent assays (Text S2; figs. S3 and S4). These results suggest that the miRNA dro-miR432, encoded in the W-specific region of chromosome 11, negatively regulates *PGL3* expression.

## Evolution of the ZZ/ZW system

To infer the evolutionary history of the ZZ/ZW system with miRNA-mediated regulation within the genus *Dioscorea*, we performed synteny analysis between white Guinea yam and two other *Dioscorea* species, *D. tokoro* and *D. alata*, both of which harbor the XY/XX system. We previously identified sex-determination regions (SDRs) on chromosome 3 of the *D. tokoro* genome. These SDRs include the Y-specific genes *BLH9* and *HSP90*, which likely control sex determination (*11*). In *D. alata*, chromosome 6 was previously shown to contain SDRs (*12*, *31*). Synteny analysis showed that the SDR of *D. tokoro* is syntenic with the equivalent region on the sex chromosome (chromosome 6) of *D. alata* but not with that of white Guinea yam (chromosome 11). The W-specific region of the white Guinea yam genome is not syntenic with the sex chromosomes of either *D. tokoro* or *D. alata* but is instead syntenic with the equivalent regions of autosomes of these species (Fig. 4A). In white Guinea yam, putative homologs of *BLH9* and *HSP90* (proposed sex-determination genes in *D. tokoro*) are located on autosomes (chromosomes 18 and 3, respectively) and their expression is not sex-specific (figs. S5 and S6). By contrast, a homolog of *PGL3*, a proposed target gene of dro-miR432 in white Guinea yam, did not exhibit sex-specific expression in *D. tokoro* (fig. S7). These results suggest that distinct sex-determination genes were adopted within the genus, as represented by white Guinea yam and *D. tokoro*, concomitant with the evolution of different heterogametic sex determination systems, i.e., XY/XX and ZZ/ZW. Rearrangements of sex chromosomes and the establishment of different heterogametic systems have been reported in dioecious flowering plants, with representative examples found in the genera *Populus* and *Salix* in the Salicaceae family. Sex determination mediated by an ortholog of the *Arabidopsis* pair RESPONSE REGULATOR 16/17 (ARR16/17) is conserved in these genera (*32–34*). By contrast, the two *Dioscorea* species have adopted distinct sex-determination genes in different heterogametic systems, highlighting the diverse evolution of sex-determination systems in plants.

**Fig. 4.**
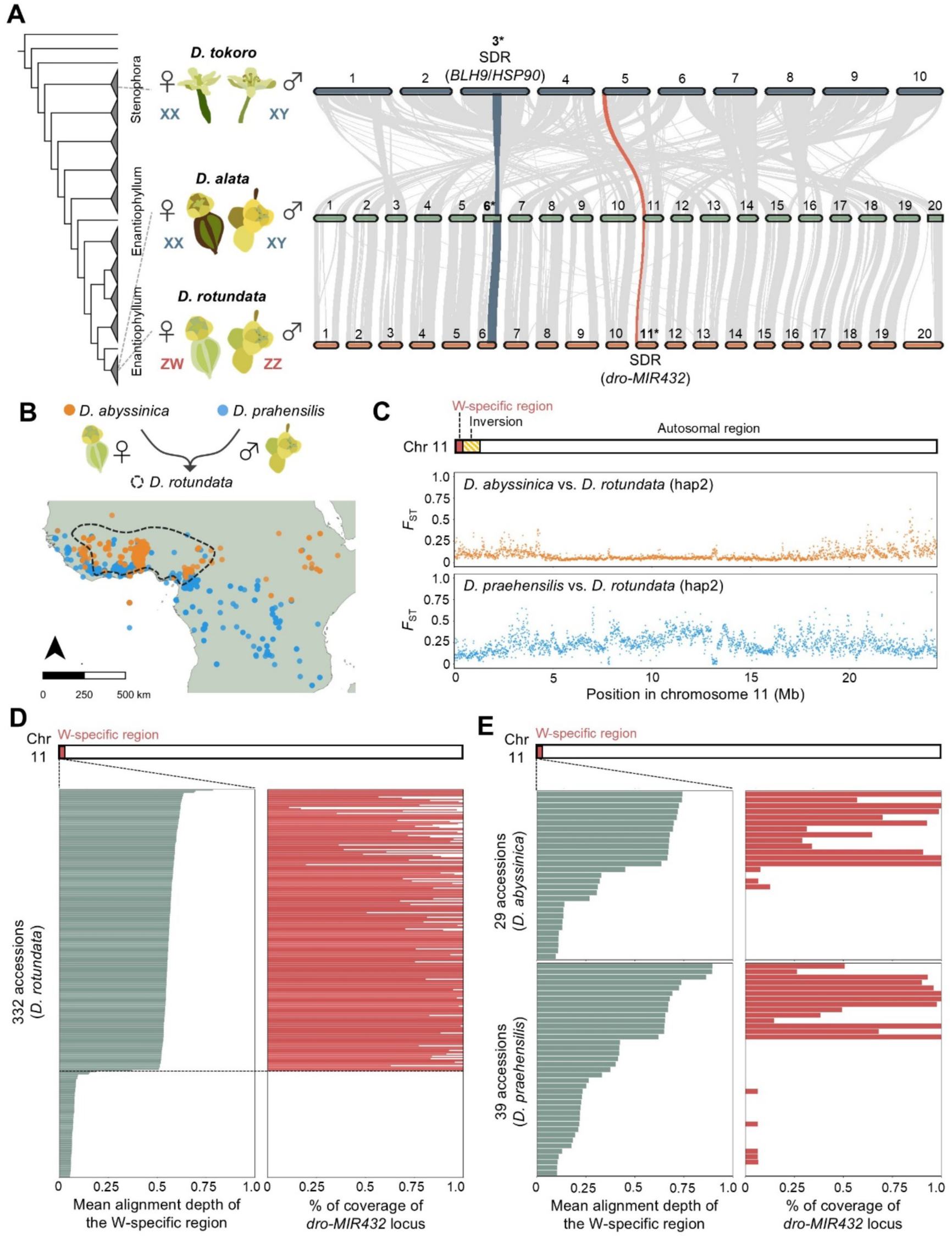
Evolution of the ZZ/ZW system with the W-specific region within the genus *Dioscorea.* **(A)** Gene order–based synteny analysis of *Dioscorea tokoro*, *D. alata*, and *D. rotundata* haplotype 2 chromosomes, suggesting that distinct sex-determination genes or miRNAs were established within the genus. Blue bands between *D. tokoro* and *D. alata* indicate synteny blocks in the sex determination regions (SDRs) on chromosome 3 (Y chromosome) in *D. tokoro* and their corresponding regions in *D. alata*. Red bands indicate synteny blocks in the W-specific region of chromosome 11 in *D. rotundata* and the corresponding regions in the other two species. Asterisks indicate the sex chromosomes in each species. **(B)** Diagram of the hybridization origin of white Guinea yam, as reported by Sugihara et al. (2020). White Guinea yam was derived from crosses between the wild savannah–adapted species *Dioscorea abyssinica* (maternal ancestor) and the rainforest species *D. praehensilis* (paternal ancestor). The lower panel shows the geographical distributions of white Guinea yam and its parental species. **(C)** Distribution of fixation index (*FST*) values across the W chromosome. **(D)** Mean alignment depth of the W-specific region and percentage of coverage of the *dro-MIR432* locus obtained from Illumina short-read sequences from 332 white Guinea yam accessions. The coverage percentage of the *dro-MIR432* locus was defined as the proportion of nucleotide positions within the locus with a normalized alignment depth > 0.2. **(E)** Mean alignment depth of the W-specific region and percentage of coverage of *dro-MIR432* locus calculated from Illumina short-read sequences from 29 accessions of the maternal ancestor *D. abyssinica* and 39 accessions of the paternal ancestor *D. praehensilis*.

To address the timing of the establishment of the ZZ/ZW system relative to the XY/XX system, we calculated the distribution of synonymous site divergence (*dS*) between the putative orthologous genes located in the inverted regions of the Z and W chromosomes (fig. S8A). We identified 158 genes in the inverted region on the W chromosome that had corresponding genes on the Z chromosome; these genes were defined as Z–W gametologs. The median *dS* value of the 158 Z–W gametologs was 0.024, which is comparable to the median intraspecific *dS* value between all orthologous or paralogous gene pairs of the haplotype 1 and 2 assemblies for *D. rotundata* (0.033). The median *dS* value of Z–W gametologs in the inverted region was lower than the median genome-wide divergence between white Guinea yam and *D. alata* (median *dS* value: 0.10) or white Guinea yam and *D. tokoro* (median *dS* value: 0.49) (fig. S8B), suggesting that the divergence of the inverted region between the Z and W chromosomes occurred more recently than the divergence between white Guinea yam and the two other *Dioscorea* species with the XY/XX system.

Next, we examined the origin of the W-specific region and dro-miR432 during the hybridization process that yielded white Guinea yam. We previously (*13*) showed that white Guinea yam originated from crosses between the wild savannah–adapted species *Dioscorea abyssinica* (maternal origin) and the rainforest species *D. praehensilis* (paternal origin). (Fig. 4B). The fixation index (*FST*) between white Guinea yam and its two wild ancestral species pointed to a greater genomic contribution from the maternal ancestor *D. abyssinica* to the sex chromosome of white Guinea yam (*13*). To evaluate the genetic distances between white Guinea yam and the two ancestral species along the W-specific regions, we calculated *FST* values across the W chromosome using the newly obtained genome assembly from this study. The distribution of *FST* values across the W chromosome indicated that a large proportion of autosomal regions showed a shorter genetic distance to the maternal ancestral species *D. abyssinica* than to the paternal ancestral species *D. praehensilis*, as evidenced by very low *F*ST values (Fig. 4C), consistent with the previous study (*13*). Following the hybridization between *D. abyssinica* and *D. praehensilis*, the corresponding *D. abyssinica-*derived regions likely became fixed in the autosomal regions of the white Guinea yam sex chromosomes. In contrast to the autosomal region, the W-specific region and the adjacent inverted region in the white Guinea yam genome show similar genetic distances to both parental species (Fig. 4C), suggesting that the genome sequences derived from both parental species might be preserved in the W-specific region in white Guinea yam.

To infer the presence of the W-specific region in the two parental species, we performed coverage analysis of Illumina short-read sequences from 332 accessions of white Guinea yam, 29 accessions of *D. abyssinica*, and 39 accessions of *D. praehensilis.* The 332 white Guinea yam accessions, maintained at the International Institute of Tropical Agriculture (IITA) in Nigeria, represent the genetic diversity of landraces and improved lines cultivated across West Africa (*13*). The mean alignment depth of the W-specific region showed two distinct patterns among the 332 white Guinea yam accessions, with approximately 0.5× depth or no (0×) depth (Fig. 4D). The two distinct patterns likely represent accessions with a ZW and ZZ genotype, respectively. All 241 accessions with the putative ZW genotype showed sequence coverage at *dro-MIR432* locus, whereas all 91 accessions with the putative ZZ genotype showed no coverage at this locus (Fig. 4D). These results suggest that diverse accessions of white Guinea yam have retained the ZZ/ZW system with dro-miR432.

We applied the same approach to the two ancestral species. The 29 *D. abyssinica* accessions and 39 *D. praehensilis* accessions showed variation in the mean alignment depth of the W-specific region. Accessions with greater alignment depth in the W-specific region also tended to show coverage at *dro-MIR432* locus (Fig. 4E), suggesting that the two ancestral species likely retained the ZZ/ZW system with the W-specific regions carrying dro-miR432. To test whether the W-specific sequences arose independently in each parental species or were already present before the two species diverged, we conducted phylogenetic analysis of the W-specific regions from white Guinea yam, *D. abyssinica*, and *D. praehensilis* using *D. alata* as an outgroup. In the resulting phylogenetic tree, the W-specific regions from *D. abyssinica* and *D. praehensilis* did not form clearly separated clades, suggesting that the W-specific region may have originated prior to the speciation of the two parental species of white Guinea yam (fig. S9).

In summary, the unstable sex expression observed in ZW individuals of white Guinea yam likely results from miRNA-mediated regulation of the ZZ/ZW system. This regulatory mechanism appears to have been established more recently than the divergence between the lineage of white Guinea yam and the lineage of *D. alata* with an XY/XX system. The extent of dro-miR432-mediated repression of *PGL3* appears to be variable. We speculate that *dro*-*MIR432* expression, pri*-*miR432 processing, or dro-miR432-mediated targeting of *PGL3* are affected by environmental factors. Further studies of the molecular mechanism underlying the variation in dro-miR432-mediated suppression of *PGL3* might enable the artificial control of dro-miR432 activity, which may allow stable monoecious accessions to be generated, promoting the fixation of desirable traits through self-pollination. Moreover, examining the evolutionary consequences of flexible floral sex determination in white Guinea yam should improve our understanding of sex determination in plants.

## Acknowledgments

Acknowledgments follow the references and notes list but are not numbered. Start with text that acknowledges non-author contributions (including disclosure of any editing services or AI-assisted technologies used in preparation of the manuscript), then complete each of the sections below as separate paragraphs.

## Author contributions

Conceptualization: RT, AA, KI, RM, AA, SN, MS, SY Resources: KI, RM, SY, PA, PA, AA

Data curation: KI, RM, SN, PA, PA, AA Formal analysis: AK, YS, AA, SN, TS, MS

Investigation: AK, KM, KO, KO, KI, HA, KH

Project administration: RT, AA

Writing – original draft: AK, RT

Writing – review & editing: AK, RT, AA, and all authors

## Data, code, and materials availability

All data are available in the Supplementary data and public databases (NCBI: https://www.ncbi.nlm.nih.gov/bioproject/PRJNA1503423, Zenodo: https://doi.org/10.5281/zenodo.21927726)

## Supplementary Materials

## Materials and Methods

### Confirmation of sex-linked regions

#### Plant materials

The 156 F1 progeny derived from a cross between the monoecious parent (TDr04/219) and male parent (TDr97/777) (*1*) were used to confirm the sex-linked region. The 156 F1 progeny comprised 40 female or monoecious plants, 104 male plants, and 12 non-flowering plants.

#### Genome-wide association study

To identify genomic regions significantly associated with sex phenotypes, genome-wide association study (GWAS) was performed. The Illumina short-read sequences of the F1 progeny were aligned to the consensus genome assembly (*1*). A total of 1,098,202 single nucleotide polymorphism (SNP) markers were obtained using Samtools v1.8 with the mpileup command with the option “-t DP,AD,SP,ADF,ADR,INFO/ADF,INFO/ADR -B -Q 18 -C 50”. GWAS was performed based on the SNP markers and sex phenotypes of the 156 F1 individuals using the R package rrBLUP v4.6 with the GWAS command with the option “K=NULL, n.PC=0, min.MAF=0.05”. The log10-transformed *p*-values (–log10(*p*)) were visualized as Manhattan plots. The false discovery rate (FDR) was set to 0.05 and was corrected by Bonferroni correction.

### Genome assembly and gene annotation

#### Plant materials

The *D. rotundata* TDr96_F1 reference genome, which was generated in previous studies (*1*, *2*), was used for haplotype-resolved genome assembly. TDr96_F1 is diploid (2n = 2× = 40) and is presumed to be ZW (*2*).

#### *De novo* assembly

Genomic DNA was extracted from fresh leaves of TDr96_F1, and PacBio long-read sequencing was conducted at Cornell University. Omni-C chromatin conformation capture data from TDr96_F1 were also generated at Iwate Biotechnology Research Center (Table S6). The PacBio long reads were assembled using Hifiasm v0.19.8-r603 (*3*) in Hi-C integration mode using the Omni-C chromatin conformation capture data. This step generated three assemblies: 1,122 primary contigs with an N50 value of 24,924,176 bp and a total size of 652.2 Mb; 1,209 haplotype 1 contigs with an N50 value of 17,927,676 bp and a total size of 630.6 Mb; and 280 haplotype 2 contigs with an N50 value of 15,137,522 bp and a total size of 570.4 Mb. The primary contigs, pseudo-haploid contigs, appeared to be the most complete, with 14 telomere-to-telomere contigs. The haplotype 1 and 2 assemblies include eight and nine telomere-to-telomere contigs, respectively.

#### Homology-based scaffolding

Chromosomal-scale genome sequences were generated from the assembled contigs by homology-based scaffolding onto the pseudochromosomes (Assembly Ver. 2), which were constructed in (*1*) (fig. S1A). First, primary contigs, representing the most complete assembly, were scaffolded to the pseudochromosomes (Ver. 2) using RagTag v2.1.0 (*4*) with the option “scaffold”. A comparison between the newly obtained pseudochromosomes and the previously obtained pseudochromosomes (Ver. 2) suggested that the newly obtained pseudochromosomes were scaffolded without major deletions (fig. S1B). A dot plot was generated using D-Genies (*5*). Based on the new pseudochromosomes obtained from the primary contigs, the haplotype 1 and 2 assemblies were scaffolded using RagTag v2.1.0 (*4*) with the option “scaffold”. After obtaining pseudochromosomes from the haplotype 1 and 2 assemblies, the contig orientations in each chromosome were corrected based on the telomere repeats. The telomere repeats were identified using quarTeT v1.2.5 (*6*) with the option “TeloExplorer -c plant” (Table S7). To evaluate the completeness of the final assemblies, BUSCO (Benchmarking Universal Single-Copy Orthologs) v5.2.2 (*7*) was utilized with “genome” as the assessment mode and Embryophyta odb10 as the database (Table S1). The collinearity between pseudochromosomes from haplotypes 1 and 2 was confirmed using a dot plot generated with D-Genies (*5*) (fig. S1C).

#### Gene annotation

Previously predicted genes in the reference genome Ver. 2 (*1*) were lifted over to the haplotype 1 and 2 assemblies using the liftOver function in ALLMAPS (*8*). Among the 35,498 genes with 66,561 transcript variants in the reference genome Ver. 2, 32,881 genes with 63,386 transcript variants were lifted over to the haplotype 1 assembly, and 32,908 genes with 63,487 transcript variants were lifted over to the haplotype 2 assembly (Table S1). Finally, gene order–based synteny of the haplotype 1 and 2 reference genomes was detected using MCscan (*9*). The orthologous regions were identified using the “jcvi.compara.catalog ortholog” function in JCVI (*9*) with the option “--cscore=.99” and the “jcvi.compara.synteny screen” function with at least 50 collinear gene blocks. The detected collinearity was visualized using the “jcvi.graphics.karyotype” function (fig. S1D).

### Identification of the W-specific region

#### Alignment-based dot plot analysis

An alignment-based dot plot of chromosome 11 between the haplotype 1 and haplotype 2 assemblies was generated using D-Genies (*5*).

#### Analysis of short-read coverage

To characterize the haplotype 2-specific region, Illumina short reads of the female individual TDr97_00917 and male individual TDr97_00777 were obtained (*2*). For quality control of the Illumina short reads, adapters and reads of <50 bp, as well as low-quality reads with an average read quality score <20, were removed using FaQCs v2.10 (*10*). The filtered short reads were aligned to the haplotype 1 and 2 reference genome assemblies. Sequence alignment was conducted using BWA v0.7.17-r1188 (*11*) with the BWA-MEM algorithm setting. The alignment depths were calculated with Samtools v1.16.1 (*12*); in this step, only reads with mapping quality ≥ 40 were counted. The alignment depths were normalized by dividing by the mean depth of all positions on the chromosomes. To reduce the effect of error on the depth, a sliding window approach was employed (window size = 150 kb, step size = 10 kb). Regions were identified in which the female depth was 0.5 (half of mean normalized depth) ± 0.15 and the male depth was < 0.15.

### Identification of candidate genes for sex determination

#### Identification of highly expressed genes in female flowers

Differential gene expression analysis was performed using the filtered RNA-seq data from male flowers, female flowers, and non-reproductive organs. Non-reproductive organs contained 11 samples: axillary bud, leaf, petiole, pulvinus, rachis, stem, spine, root, tuber head, tuber middle, and tuber tail. The trimmed RNA-seq reads obtained from (*1*) were aligned to the haplotype 2 assembly using HISAT v2.2.1 (*13*). The mapped reads were counted using the featureCounts function of Subread v2.0.1 (*14*). The minimum fragment length was set to 19, and the attribute type was set to transcript id. Differential gene expression analysis was performed using DESeq2 v3.15 (*15*) in R v4.1.1 for two comparisons: male vs. female flowers at three stages of development and male flowers at three stages of development vs. non-reproductive organs. The false discovery rate threshold was set to 0.05.

#### Identification of candidate genes for sex determination

The W-specific region contained 17 potential protein-coding genes. Of these genes, 12 were highly expressed in all comparisons of the three stages of female vs. male flowers based on the hypothesis test for differential expression of the RNA-seq data using negative binomial generalized linear models (*p*-value < 0.05). To narrow down the list of candidate genes, the expression levels of the 12 genes were compared between female flowers and non-reproductive organs, but no gene showed differential expression in this comparison.

### Identification of candidate miRNAs for sex determination

#### Plant materials

Small RNA-seq data were obtained from female and male *D. rotundata* flowers collected at three developmental stages (stage 1–3). Female flowers were derived from three accessions (DrDRS002, DrDRS027, and DrDRS028). Male flowers were derived from three accessions (DrDRS020, DrDRS063, and DrDRS067). Three biological replicates were included for each accession at each developmental stage, resulting in a total of 54 samples (Table S8).

#### Small RNA library construction and sequencing

Small RNA-seq data were obtained from the 54 *D. rotundata* samples as previously described (*16*). Total RNA was extracted from the samples using an Ambion Plant RNA Isolation Aid (Ambion, Austin, TX, USA) following the manufacturer’s protocol with slight modifications. The samples were frozen, ground to a powder, mixed with 100 μL of Plant RNA Isolation Aid and 1 mL lysis solution, and thoroughly homogenized. The samples were centrifuged at 15,000 g in a microcentrifuge for 5 min at room temperature. The supernatants (containing total RNA) were transferred to new tubes. The small RNA fraction was isolated from total RNA using a mirVana miRNA Isolation kit (Ambion) following the manufacturer’s protocol. Small RNA libraries were constructed using a NEBNext Multiplex Small RNA Library Prep Set for Illumina (New England BioLabs, Ipswich, MA, USA) following the manufacturer’s protocol from the adapter ligation step to the PCR amplification step. For quality control, the PCR-amplified cDNAs were purified using DNA Clean & Concentrator-5 (Zymo Research, CA, USA) following the manufacturer’s protocol. After eluting the purified DNA, size selection was conducted using AMPure XP Beads following the protocol for the NEBNext Multiplex Small RNA Library Prep Set for Illumina (New England BioLabs). The quality and DNA amount of the libraries were assessed using a Qubit fluorometer (Invitrogen), an Agilent Bioanalyzer with an Agilent High Sensitivity DNA Kit (Agilent Technologies), and qPCR with a Library Quantification Kit (Takara Bio). The libraries were sequenced on the NovaSeq 6000 platform at Genebay, Yokohama, Japan (Table S8).

#### Sequence processing

For quality control of the small RNA-seq data, adapters and low-quality reads with an average read quality score < 20 were removed with FaQCs v2.10 (*10*). Reads shorter than 19 bp and longer than 25 bp were removed using Seqkit v2.3.0 (*17*). To predict miRNAs using miRDeep-P2 v1.1.4 (*18*), the filtered reads were preprocessed into the designated format. The filtered reads were parsed into FASTA format with Seqkit v2.3.0, and redundant sequences were removed.

#### miRNA prediction and annotation

The miRNAs were predicted as described previously (*19*) with several modifications. Novel miRNAs were detected from preprocessed reads using the miRDP2-v1.1.4_pipeline.bash script of miRDeep-P2 v1.1.4. The predicted miRNA sequences and their corresponding precursor and primary information were extracted using the perl script parse_miRDP2_prediction.pl (*19*). The positions of miRNA gene loci in the haplotype 2 assembly were determined using the BLASTN function of BLAST+ v2.2.31 (*20*).

#### Identification of highly abundant miRNAs in female flowers

The trimmed small RNA-seq reads were aligned to the predicted miRNA sequences with Bowtie v1.3.1 (*21*) with the option “-v 0 --norc”. Only small RNA-seq reads that were perfectly aligned to the sequences in the miRNA sequences were used to quantify miRNA abundance. The mapped reads were counted using the Perl script bam2ref_counts.pl, and the read counts data for each sample were combined using the Perl script combine_htseq_counts.pl (*19*). Differential expression analysis was performed using DESeq2 v3.15 (*15*) in R v4.1.1 for the comparisons of male vs. female flowers at three stages of development. The false discovery rate threshold was set to 0.05. Predicted miRNAs with zero read counts across all samples were excluded from subsequent analysis.

#### Identification of candidate miRNAs for sex determination

The W-specific region contained a miRNA gene locus, which was predicted to produce the 24-bp mature miRNA dro-miR432. dro-miR432 was highly abundant in the comparison of the three stages of female vs. male flowers based on the hypothesis test for differential expression of the small RNA-seq data using negative binomial generalized linear models (*p*-value < 0.05). The abundance of mature miRNAs was also evaluated based on the log2-normalized ratio of the mean of normalized counts (log2[fold-change], log2FC). dro-miR432 showed more than 4-fold higher abundance (log2FC > 2) during the three stages of female flowers than those from male flowers.

#### Identification of candidate target genes

Candidate target genes of dro-miR432 were predicted using psRNATarget (A Plant RNA Target Analysis Server) (*22*) with scoring schema V2 using cDNA sequences of the haplotype 2 assembly as cDNA target. The default setting of scoring schema V2 was used: maximum expectation 5, penalty for G:U pair 0.5, penalty for other mismatches 1, extra weight in seed region 1.5, seed region 2–13 NT, number of mismatches allowed in seed region 2, length for complementarity scoring (HSP size) 19, penalty for opening gap 2, penalty for extending gap 0.5, translation inhibition range 10–11 NT. Expectation values were obtained, i.e., the penalty values for the mismatches between mature small RNAs and the target sequence; the gene with the smallest expectation value was selected as the putative target gene.

#### Phylogenetic analysis of the target gene

Phylogenetic analysis of PGL3 from *D. rotundata* and its homologs was performed using the protein sequences of PGL3, BURP domain–containing proteins from *Oryza sativa*, BURP domain–containing proteins from *Arabidopsis thaliana*, and putative homologs of PGL3 from monocotyledonous species (*Asparagus officinalis*, *Dioscorea alata*, *Dioscorea tokoro*, and *Phoenix dactylifera*). The sequences of *O. sativa* BURP domain–containing proteins were obtained from NCBI based on (*23*), and the sequences of *A. thaliana* BURP domain–containing proteins were obtained from NCBI based on (*24–26*) (Table S9). The putative homologs (<1 × 10^−10^ in BLASTP analysis, max_target_seqs = 2) of PGL3 were obtained from the protein sequences of four species: *Asparagus officinalis* (NCBI RefSeq assembly; GCF_001876935.1), haplotype 1 assembly of *Dioscorea alata* (*27*), male haplotype 1 assembly of *Dioscorea tokoro* (*16*), and *Phoenix dactylifera* (NCBI RefSeq assembly; GCF_009389715.1). The sequences were aligned using MAFFT v7.475 with the L-INS-i strategy (*28*). Based on the alignment, a phylogenetic tree was reconstructed with the maximum likelihood method using IQ-Tree v2.1.2 (*29*). The bootstrap values were calculated using ultrafast bootstrap approximation with 1,000 replications. The tree was visualized using Interactive Tree Of Life (iTOL) v7.0 (*30*). The AtRD29A protein encoded by At5g52310 was used as the outgroup.

### Dual-luciferase reporter assay

miRNA activity was quantitatively evaluated using a dual-luciferase assay via *Agrobacterium tumefaciens*–mediated infiltration of *Nicotiana benthamiana* leaves as previously described (*33*) with some modifications.

#### Plasmids for dual-luciferase reporter expression

Dual-luciferase reporters were constructed in the pBIB binary vector backbone (*31*), which was derived from pBIN19. As shown in Fig. 3A, the dual-luciferase reporter plasmid contains two luciferase expression cassettes: a *Renilla reniformis* luciferase expression cassette (P*nos*:*R-Luc*:T*g7*) that functions as an internal control to standardize expression, and a firefly luciferase expression cassette (P*nos*:*F-Luc*^+^:T*nos*) used to test the putative miRNA target sequence. The P*nos*:*R-Luc*:T*g7* cassette consisted of the promoter of the *Agrobacterium* nopaline synthase gene (P*nos*), the coding region of the *R. reniformis* luciferase gene (*R-Luc* derived from pRL-null, Promega), and the terminator of *Agrobacterium* gene 7 (T*g7*). The P*nos*:*F-Luc^+^*:T*nos* cassette consisted of P*nos*, the coding region of the modified firefly luciferase gene (*R-Luc^+^* derived from pSP-luc+, Promega), and the terminator of the *Agrobacterium* nopaline synthase gene (T*nos*). These two expression cassettes were positioned in opposite directions on the same plasmid (pBIB-Dual2). The nucleotide sequence of pBIB-Dual2 and information about its structure were deposited in the DNA Data Bank of Japan (DDBJ) database under accession no. LC938258. pBIB-Dual2 was constructed as follows. A 1.2-kb *Nhe*I-*Nhe*I segment of the P*nos*:*hph*:T*g7* cassette in pBIB-Hyg (34) was replaced by a synthesized 1,078-bp P*nos*(partial):*R*-*Luc*:T*g7*(partial) DNA fragment (Table S10) by In-Fusion cloning, yielding pBIB-R-Luc. A 0.35-kb P*nos* fragment was amplified from pBIB-Hyg using the primer set Pnos(P)-F and Pnos(N)-R (Table S10), and a 4.3-kb *Pvu*II-*Nco*I segment of P*FT* (promoter sequence of the *Arabidopsis FLOWERING LOCUS T* [*FT*] gene) in pSP/FT:LUC^+^ (K. Onai, unpublished, accession no. LC937811 in DDBJ), which contains a P*FT*:*luc*^+^:T*nos* cassette, was replaced by the P*nos* fragment by In-Fusion cloning, giving pSP/nos:luc^+^. A 2.3-kb P*nos*:*luc^+^*:T*nos* cassette was amplified from pSP/*nos:luc^+^* using the primer set Pnos(H)-F and Tnos(E)-R (Table S10) and inserted into the *Hind*III-*Eco*RI site of pBIB-*R-Luc* by In-Fusion cloning, giving pBIB-Dual2. The double-stranded putative miRNA target sequence from *PGL3* was generated by annealing the oligonucleotides Sensor_seq_target_F and Sensor_seq_target_R (Table S10) and inserting the resulting double-stranded DNA molecule into the unique *Xba*I site of pBIB-Dual2 located downstream of the *F*-*luc*^+^ stop codon by In-Fusion cloning, giving pBIB-Dual2-tgt. pBIB-Dual2 and pBIB-Dual2-tgt were used as a negative control (shown as “Cont.” in fig. S5) and to test the target sequence (shown as “Target” in Fig. 3, fig. S4, and fig. S5), respectively. The nucleotide sequences of all constructs were confirmed by Sanger sequencing using primers listed in Table S10.

#### Plasmids for pri-miRNA expression

Each 430-bp synthesized DNA fragment of pri-miR432 and its mutated version (miRNA_seq_target_F and miRNA_seq_mutated_F, respectively, in Table S10) was replaced with a 1.9-kb *Xba*I-*Sac*I segment including *GUS* (coding sequence of the β-glucuronidase gene) in the binary vector pBI121 (*32*) by In-Fusion cloning, giving pBI-35S:Dr-miR432 and pBI-35S:Dr-miR432mut, respectively. In the mutated miRNA Dr-miR432mut, the order of nucleotides in the core sequence for target recognition was altered, but the original nucleotide composition was maintained. pBI121 (shown as “GUS” in Fig. 3, fig. S3, and fig. S4), pBI-35S:Dr-miR432 (shown as “miRNA” in Fig. 3, fig. S3, and fig. S4), and pBI-35S:Dr-miR432mut (shown as “Mutated” in Fig. 3 and fig. S3) were used as controls, for native miRNA testing, and for mutated miRNA testing, respectively. The nucleotide sequences of all constructs were confirmed by Sanger sequencing using primers listed in Table S10.

#### *Agrobacterium*-mediated transient expression in *Nicotiana benthamiana*

Transient gene expression in *N. benthamiana* was performed by *Agrobacterium tumefaciens*–mediated infiltration as previously described (*33*). Wild-type *N. benthamiana* plants were grown in soil at 24°C under long day (16-h light/8-h dark) conditions for approximately three weeks, and their leaves were used for transient gene expression. The light intensity during the light period was 70 μmol m^−2^ s^−1^ provided by white LEDs. Plasmids for reporter expression and pri-miRNA expression were introduced into *Agrobacterium tumefaciens* strain GV3101::pMP90 by electroporation, and transformants were selected on LB agar medium containing 50 μg/mL kanamycin, 50 μg/mL gentamicin, and 50 μg/mL rifampicin. Selected transformants were grown in liquid LB medium containing 50 μg/mL kanamycin and 50 μg/mL rifampicin for 21 h and in liquid LB medium containing 50 μg/mL kanamycin, 50 μg/mL rifampicin, and 15 μM acetosyringone for 3 h. The cells were collected by centrifugation at 12,000 rpm in a microcentrifuge for 1 min at room temperature, rinsed once with infiltration buffer (10 mM MES and 10 mM MgCl2, pH 5.6), and resuspended in infiltration buffer (10 mM MES, 10 mM MgCl2, and 150 μM acetosyringone, pH 5.6) at an OD600 = 0.5. The cell suspensions for dual-luciferase reporter expression and the suspension for *GUS* or pri-miRNA expression were mixed at a 1:9 (v/v) ratio as follows: (1) the reporter with the target sequence and the *GUS* expression construct (shown as “Target-GUS” in Fig. 3, fig. S4, and fig. S5), (2) the reporter with the target sequence and pri-miR432 expression construct (shown as “Target-miRNA” in Fig. 3, fig. S3, and fig. S4), (3) the reporter with the target sequence and the mutated pri-miR432 expression construct (shown as “Target-Mutated” in Fig. 3 and fig. S3), (4) the reporter without the target sequence and *GUS* expression construct (shown as “Cont.-GUS” in fig. S4), and (5) the reporter without the target sequence and pri-miR432 expression construct (shown as “Cont.-miRNA” in fig. S4). For all mixtures, only one pair of constructs was co-infiltrated per plant, and the plants were incubated for 48 h or 72 h under the same growth conditions.

#### Measurement of luciferase activity

After 48 h or 72 h of incubation, approximately 50 mg of *N. benthamiana* leaf tissue was collected from three plants using a hole punch, with three biological replicates. The leaf tissue was ground to a fine powder in liquid nitrogen and homogenized in 250 µL or 300 μL of Glo Lysis Buffer (Promega). Following centrifugation at 15,000 rpm for 1 min at 4°C, the supernatant was recovered, mixed with an equal volume of Glo Lysis Buffer, and used for the dual-luciferase assay using the Dual-Glo Luciferase Assay System (Promega). A 75-μL diluted sample was placed into each well of a 96-well plate (Nunc F96 MicroWell Polystyrene Plate white; ThermoFisher Scientific) and mixed with 75 μL of Dual-Glo Reagent (Promega). After 10 min, bioluminescence from firefly luciferase (F-Luc) was measured. Subsequently, 75 μL of Dual-Glo Stop & Glo Reagent (Promega) was added to each sample to quench the F-Luc signal and obtain bioluminescence from *Renilla* luciferase (R-Luc). After 10 min, R-Luc activity was measured. Bioluminescence from both F-Luc and R-Luc was measured automatically every 1 min at 15 time points at 23°C using an automated bioluminescence-monitoring apparatus (model CL96-4; Churitsu Electric Corp., Nagoya, Japan) (Data S1, S2). To calculate the F-Luc/R-Luc ratios, the average measurements from the 3–5 time points were used. For the first step of statistical analysis, bioluminescence values were corrected by subtracting the background signal, defined as the average bioluminescence of two empty wells. After background correction, the F-Luc/R-Luc ratios were calculated for each sample. Finally, relative F-Luc/R-Luc ratios were obtained by normalizing the average ratio under control conditions (co-infiltration with the *GUS* expression construct). The relative F-Luc/R-Luc ratios among three co-infiltration pairs (Target-GUS, Target-miRNA, and Target-Mutated) were compared using pairwise Welch’s *t*-tests with Holm-adjusted *p*-values. The relative F-Luc/R-Luc ratios between two co-infiltration pairs (Cont.-GUS vs. Cont.-miRNA, and Target-GUS vs. Target-miRNA) were compared using Welch’s t-test.

### Comparative genomic analysis with *D. alata* and *D. tokoro*

#### Synteny analysis

To compare the sex chromosomes of *D. rotundata*, *D. alata*, and *D. tokoro*, gene order–based synteny of the chromosomes was detected using MCscan (*9*). The orthologous regions were identified from the haplotype 2 assembly of *D. rotundata*, the haplotype 1 assembly of *D. alata* (*27*), and the male haplotype 1 assembly of *D. tokoro* (*16*) using the “jcvi.compara.catalog ortholog” and the “jcvi.compara.synteny screen” function with at least 10 collinear gene blocks. The detected collinearity was visualized using the “jcvi.graphics.karyotype” function.

#### Phylogenetic analysis of BLH9, *HSP90*, and PGL3

Phylogenetic analysis of BLH9 was performed using protein sequences of TALE superfamily proteins from *A. thaliana* and its putative homologs from *D. rotundata* and *D. tokoro.* The sequences of *A. thaliana* TALE superfamily proteins were obtained from (*16*). The putative homologs (<1 × 10^−10^ in BLASTP analysis, max_target_seqs = 20) of TALE superfamily proteins were obtained from the protein sequences of haplotype 2 of *D. rotundata* and male haplotype 1 of *D. tokoro* (*16*). Phylogenetic analysis of *HSP90* was performed using the mRNA sequences of HSP90 family proteins from *A. thaliana* and its putative homologs from *D. rotundata* and *D. tokoro.* The sequences of *A. thaliana* HSP90 family proteins were obtained from (*16*). The putative homologs (<1 × 10^−10^ in TBLASTN analysis, max_target_seqs = 20) of *HSP90* family genes were obtained from the mRNA sequences of haplotype 2 of *D. rotundata* and male haplotype 1 of *D. tokoro.* Phylogenetic analysis of PGL3 was performed using the protein sequences of BURP domain–containing proteins from *A. thaliana* and its putative homologs from *D. rotundata* and *D. tokoro.* The sequences of *A. thaliana* BURP domain–containing proteins were obtained from NCBI (Table S9). The putative homologs (<1 × 10^−10^ in BLASTP analysis, max_target_seqs = 20) of BURP domain–containing proteins were obtained from the protein sequences of haplotype 2 of *D. rotundata* and male haplotype 1 of *D. tokoro*. In each phylogenetic analysis, the sequences were aligned using MAFFT v7.475 with the L-INS-i strategy (*28*). Based on the alignments, a phylogenetic tree was reconstructed with the maximum likelihood method using IQ-Tree v2.1.2 (*29*). Bootstrap values were calculated by ultrafast bootstrap approximation with 1,000 replications. The trees were visualized using Interactive Tree Of Life (iTOL) v7.0 (*30*).

#### Transcriptomic comparison of *BLH9*, *HSP90*, and *PGL3*

Differential gene expression analysis was performed using the filtered RNA-seq data from male flowers, female flowers, and non-reproductive organs of *D. rotundata* and *D. tokoro*. The trimmed RNA-seq reads for *D. rotundata* and *D. tokoro* were obtained from (*1*) and (*16*), respectively. The RNA-seq reads were aligned to the haplotype 2 assembly of *D. rotundata* or the male haplotype 1 assembly of *D. tokoro* using HISAT v2.2.1 (*13*). The mapped reads were counted with the featureCounts function in Subread v2.0.1 (*14*). The minimum fragment length was set to 19, and the attribute type was set to transcript id.

#### Synonymous divergence (*dS*) analysis

To identify Z-linked and W-linked genes located in the inverted region, gene order–based synteny of the Z and W chromosomes was detected using MCscan (*9*). The orthologous regions were identified using the “jcvi.compara.catalog ortholog” function with the option “--cscore=.99” and the “jcvi.compara.synteny screen” function with at least 20 collinear gene blocks. Synonymous divergence (*dS*) between Z- and W-linked gametologs was obtained as previously described (*34–36*) with slight modifications. Orthologous or paralogous gene pairs (<1 × 10^−50^ in BLASTP analysis, max_target_seqs = 2) were identified from three comparisons: haplotype 1 and 2 of *D. rotundata*, haplotype 2 of *D. rotundata* and haplotype 1 of *D. alata* (*27*), and haplotype 2 of *D. rotundata* and male haplotype 1 of *D. tokoro* (*16*). The gene pairs were aligned in codon frames using MAFFT v7.475 (*37*) with the option “--localpair --maxiterate 1000” and Pal2Nal v14 (*38*). Based on the in-codon-frame alignments, the Jukes and Cantor corrected *dS* values were calculated using KaKs_Calculator v2.0 (*39*). All predicted genes (32,671 genes) with complete coding sequences from the *D. rotundata* haplotype 2 assembly were used as references. In total, 51,529 orthologous or paralogous gene pairs were identified by comparison to haplotype 1 of *D. rotundata*, 36,014 gene pairs by comparison to haplotype 1 of *D. alata*, and 59,863 gene pairs by comparison to male haplotype 1 of *D. tokoro*. A total of 158 genes in the inverted region on the W chromosome had corresponding genes in the Z chromosome and were therefore defined as Z–W gametologs. The comparison and distributions of *dS* values were visualized using R v4.4.1.

### Comparative genome analysis with the parental species *D. abyssinica* and *D. praehensilis*

#### Plant materials

Illumina short reads from 332 *D. rotundata* accessions (Data S3), 29 *D. abyssinica* accessions, 39 *D. praehensilis* accessions and two accessions of *D. alata* (Data S4) were obtained from (*1*).

#### Analysis of fixation inde

For quality control of the Illumina short reads, adapters and reads <50 bp, as well as low-quality reads with an average read quality score <20, were removed using FaQCs v2.10 (*10*). The filtered short reads were aligned to the haplotype 1 and 2 reference genomes. Sequence alignment was conducted using BWA v0.7.17-r1188 (*11*) with the BWA-MEM algorithm setting. Based on these alignments, variant calling was conducted using Bcftools v1.15 (*12*) with the following commands: (i) mpileup command; (ii) call command with the option “-v - m”; (iii) view command with “-v snps”; (iv) filter command with the options “-i ‘QUAL>30 && DP>5’”. *F*ST statistics were calculated using VCFtools v0.1.16 with the option “--fst-window-size 10000” for comparisons between *D. rotundata* and *D. abyssinica*, and between *D. rotundata* and *D. praehensilis*.

#### Analysis of short-read coverage of the W-specific region

The alignment depths were calculated with Samtools v1.16.1 (*12*); in this step, reads with mapping quality ≥ 40 were counted. The alignment depths were normalized by dividing by the mean depth of all positions on the chromosomes. Following normalization, the mean alignment depth of the W-specific region and the coverage percentage of *dro-MIR432* locus were calculated for each accession. The coverage percentage of the *dro-MIR432* locus was defined as the proportion of nucleotide positions within the locus with a normalized alignment depth > 0.2.

#### Phylogenetic analysis of the W-specific region

Phylogenetic analysis was performed using nucleotide sequences of the W-specific regions from *D. rotundata*, *D. abyssinica*, and *D. praehensilis* accessions. After aligning to the haplotype 2 assembly, accessions with normalized alignment depths greater than 0.2 at ≥75% of sites in the W-specific region were considered to have sufficient coverage and were included in the phylogenetic analysis. In total, 219 *D. rotundata* accessions, seven *D. abyssinica* accessions, and eight *D. praehensilis* accessions were selected. The sequences were aligned using MAFFT v7.475 with the L-INS-i strategy (*28*). Based on the alignment, a phylogenetic tree was reconstructed with the maximum likelihood method using IQ-Tree v2.1.2 (*29*). The bootstrap values were calculated by ultrafast bootstrap approximation with 1,000 replications. The tree was visualized using Interactive Tree Of Life (iTOL) v7.0 (*30*). *D. alata* was used as the outgroup.

## Supplementary Text S1

### Haplotype-resolved genome assemblies of *Dioscorea rotundata*

A haplotype-resolved genome assembly from a monoecious individual of *D. rotundata* (TDr96_F1) was constructed using PacBio HiFi whole-genome sequencing reads and Omni-C chromatin conformation capture data. The resulting haplotype 1 assembly consisted of 1,209 contigs (630.6 Mb in total), while the haplotype 2 assembly comprised 280 contigs (570.4 Mb in total). The haplotype 1 and 2 assemblies contained eight and nine telomere-to-telomere contigs, respectively. BUSCO (*7*) analysis showed that the percentages of complete (single-copy and duplicated) embryophyte BUSCOs were 98.3% for the haplotype 1 assembly and 98.5% for the haplotype 2 assembly (Table S1). The assembled contigs were scaffolded onto the pseudochromosomes of the reference genome Ver. 2 (*1*), and the orientation of each contig was determined by placing the telomeric motif at the ends of each pseudochromosome. A phased pseudochromosome genome assembly Ver. 3 was obtained, consisting of 14 and 16 telomere-to-telomere pseudochromosomes for haplotype 1 and 2, respectively. All annotations in the previous assembly (*1*) were lifted over to the new assemblies (Table S1). Chromosomal synteny analysis based on collinear blocks with predicted genes revealed high collinearity between haplotypes 1 and 2 (fig. S1D). In particular, chromosome 11, which was previously reported as a sex chromosome, showed high collinearity between the two haplotypes, and both assemblies contained telomeric repeats at both chromosome ends. These results indicate that the phased and chromosome-scale genome assemblies are useful for inferring SDRs in *D. rotundata*.

## Supplementary Text S2

### Functional relevance of dro-miR432 and the target gene *PGL3*

We performed a dual-luciferase reporter assay to validate the functional relevance of the miRNA dro-miR432 and the target gene *PGL3.* We designed a dual-luciferase reporter plasmid containing the predicted target sequence (Fig. 3A) and co-infiltrated *N. benthamiana* leaves with this reporter and the pri-miRNA expression plasmid (Fig. 3B). To evaluate the candidate miRNA target, we used three types of plasmids for co-infiltration: the β-glucuronidase gene as a control (*GUS*), the native sequence of pri*-*miR432 (miRNA; Fig. 3C), and a mutated sequence of pri-miR432 (Mutated; Fig. 3D) that maintained the nucleotide composition of the native target site but had a randomized DNA sequence. *N. benthamiana* leaves were co-infiltrated with the dual-luciferase reporter (Target) and the expression construct Target-GUS, Target-miRNA, or Target-Mutated (Fig. 3E). In the assay, the presence of the miRNA resulted in significantly lower relative F-Luc activity, whereas no repression was observed with the two other co-infiltration pairs. Similar results were obtained in two independent assays (fig. S3). To further evaluate candidate miRNA targeting, we used a dual-luciferase reporter with the native *PGL3* target site inserted into the *F-Luc* expression cassette (Target) and a reporter without an insertion as the control (Control). The dual-luciferase reporter (Control) was co-infiltrated individually with either of the two expression constructs, Cont.-GUS and Cont.-miRNA. The dual-luciferase reporter (Target) was co-infiltrated individually with two expression constructs, Target-GUS and Target-miRNA (fig. S4A). In the assay, the presence of the miRNA did not repress relative F-Luc activity from the control dual-luciferase reporter, whereas it significantly repressed relative F-Luc activity from the target dual-luciferase reporter (fig. S4B, C).fig. S1.

---

**fig. S1.**
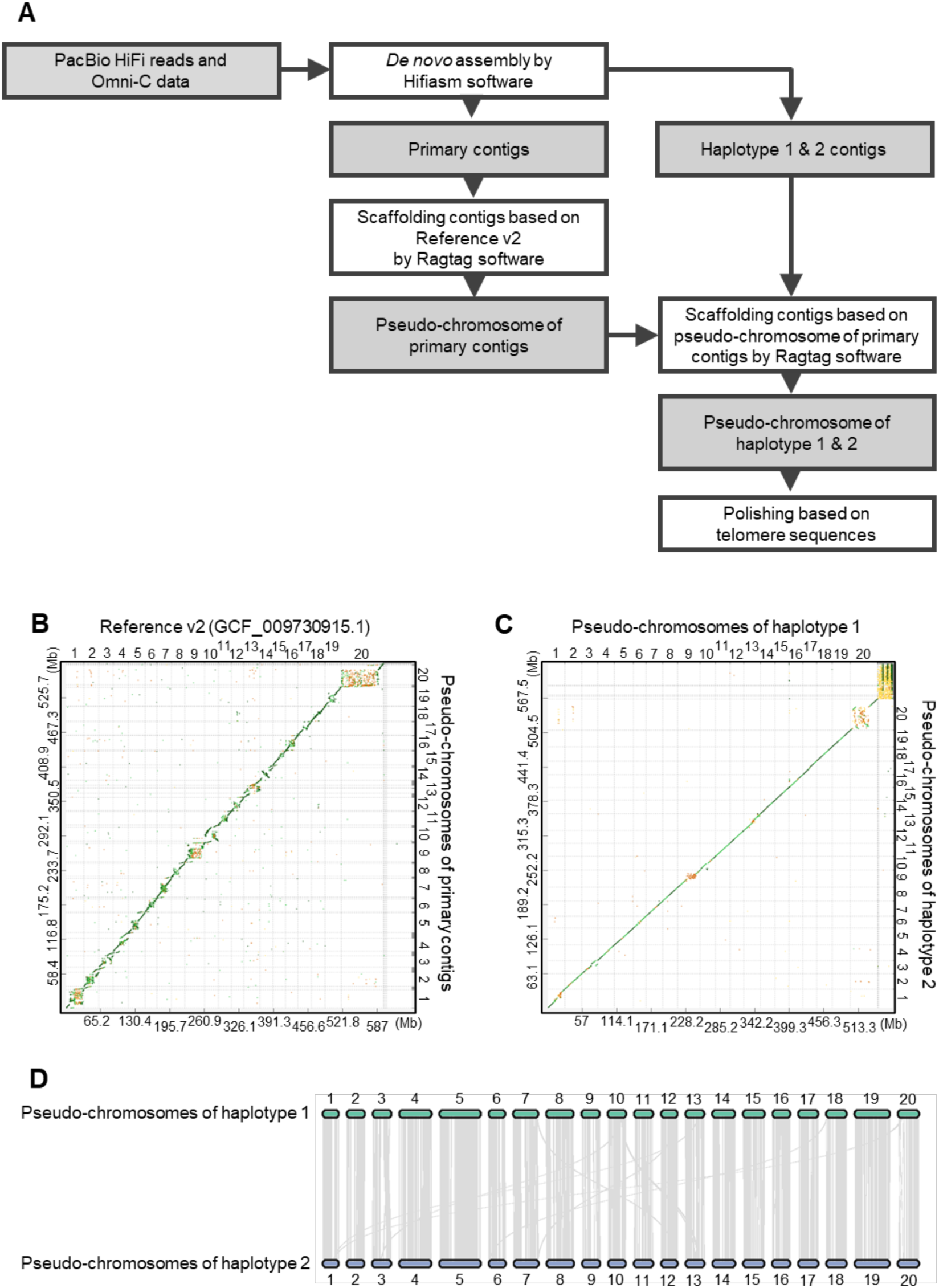
Reconstructing phased genome assemblies of *Dioscorea rotundata*. **(A)** Illustration of the pipeline used for genome assembly and homology-based scaffolding. **(B)** Alignment-based dot plots comparing the assemblies of reference Ver. 2 (*1*) and the newly constructed pseudochromosomes formed by the primary contigs. **(C)** Alignment-based dot plots comparing the pseudochromosomes of the haplotype 1 and 2 assemblies. **(D)** Chromosomal synteny analysis showing high collinearity between the haplotype 1 and 2 assemblie

**Fig. S2.**
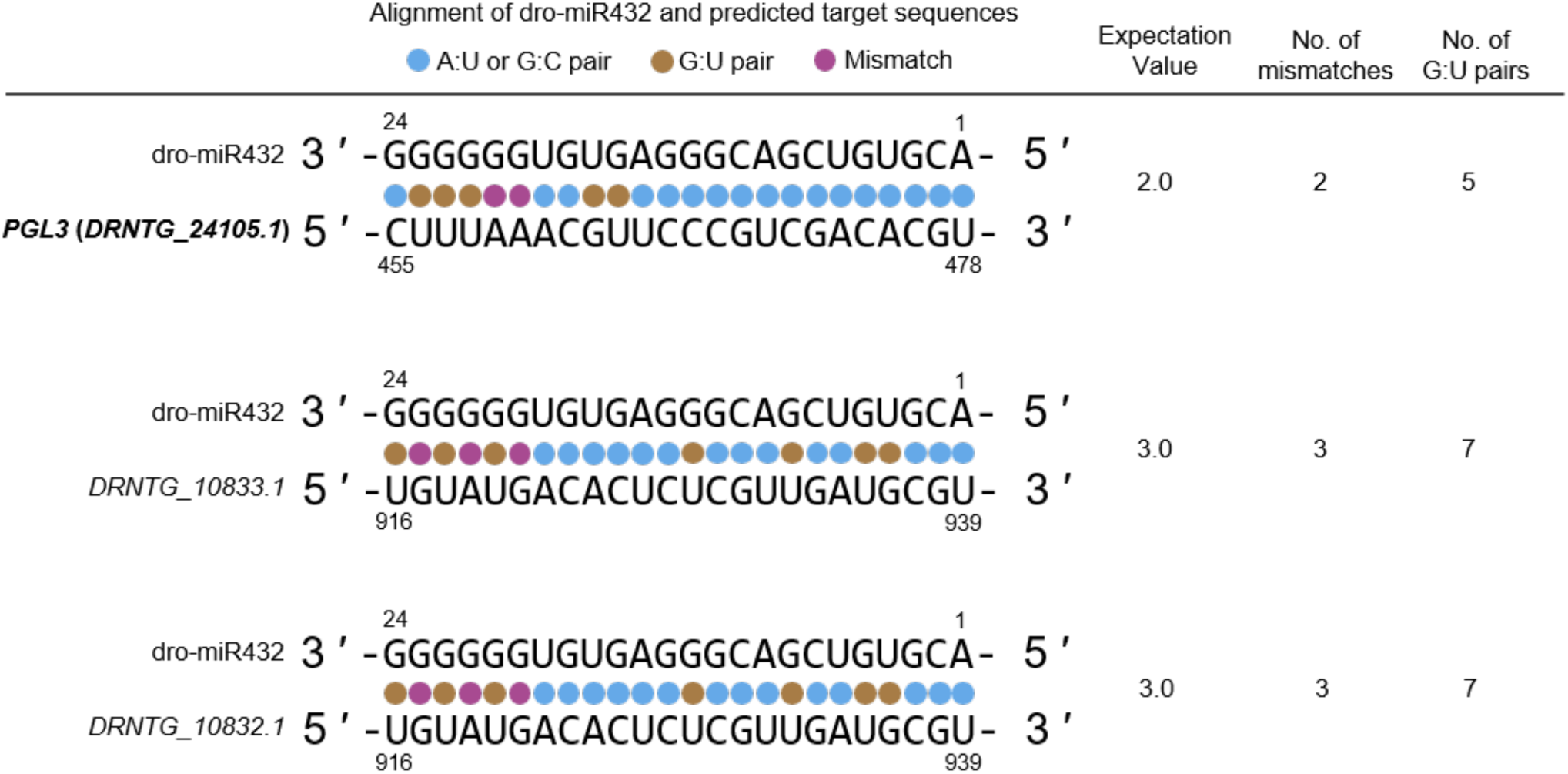
Top hits of the alignments of dro-miR432 and target sequences predicted by psRNATarget (a plant small RNA target analysis server). Candidate target genes and target sequences of dro-miR432 predicted using psRNATarget with scoring schema V2 using cDNA sequences of the haplotype 1 and 2 assemblies as cDNA targets. Expectation value indicates the penalty score for mismatches between a mature miRNA and its target sequence. Higher values indicate less similarity between the miRNA and the target candidate sequence; candidate targets with values below 3.0–5.0 are considered to be reliable in psRNATarget.

**Fig. S3.**
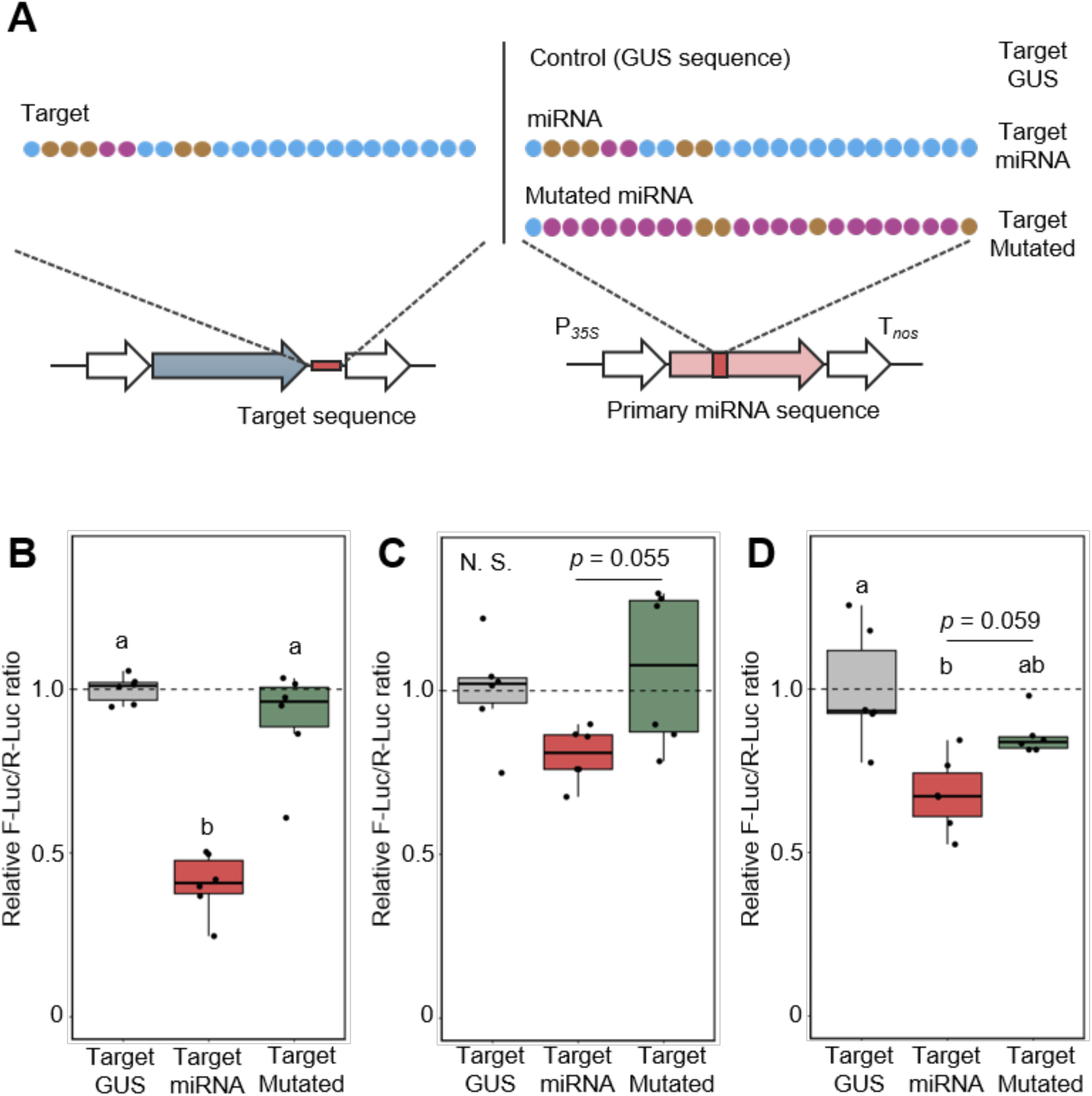
The candidate miRNA dro-miR432 targets the predicted binding sequence from *PGL3* in a dual-luciferase reporter assay in *Nicotiana benthamiana* leaves across three independent experiments. **(A)** Diagram of co-infiltrated sequences in the transient *Agrobacterium tumefaciens*–mediated infiltration assay. **(B–D)** Relative F-Luc/R-Luc ratios from three independent experiments after 72 h of incubation for the three types of co-infiltrations. N.S., not significant. The box boundaries represent the first (Q1) and third (Q3) quartiles, the horizontal line within each box indicates the median, and the whiskers extend to the most extreme data points within 1.5 × the interquartile range (IQR).

**Fig. S4.**
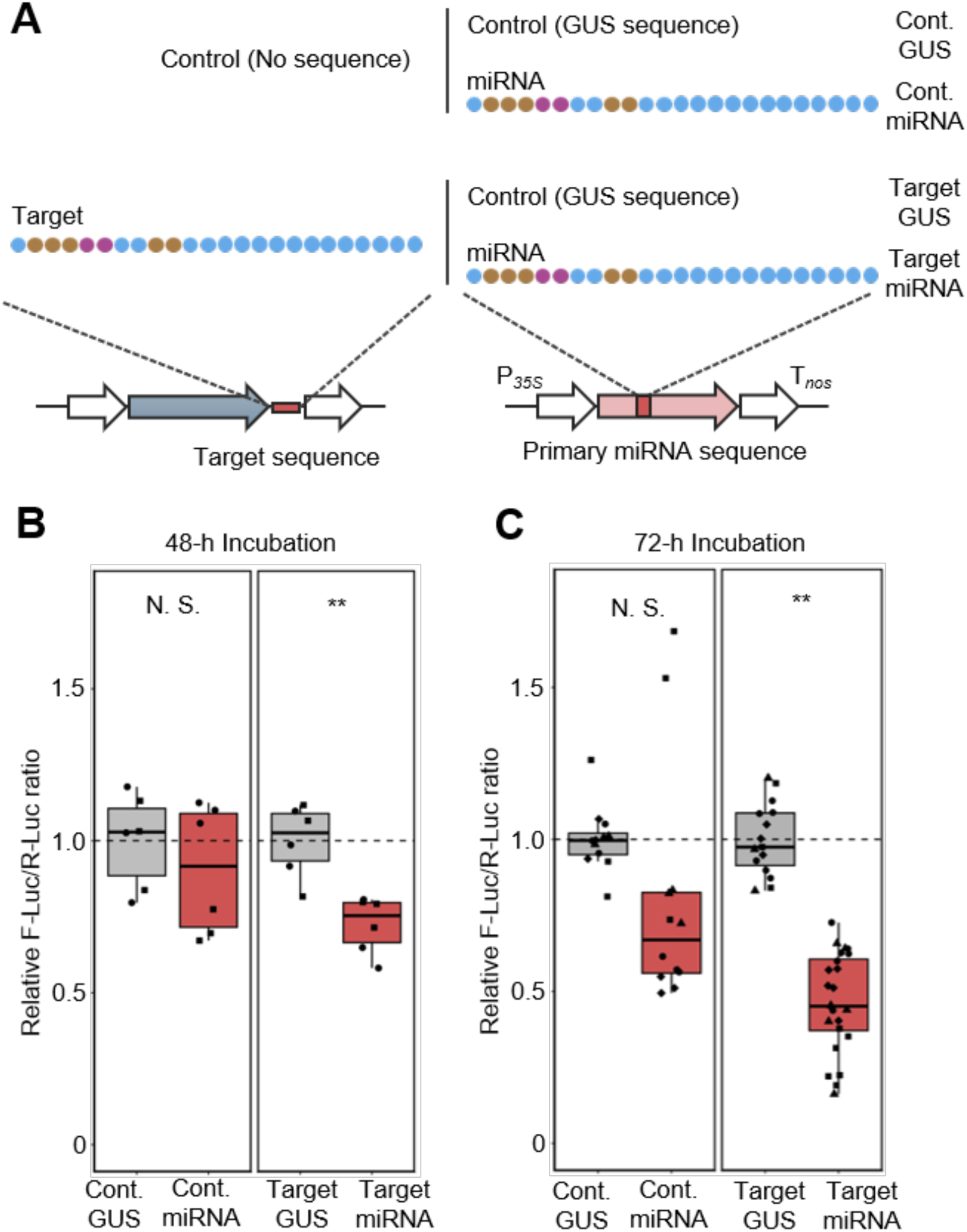
Relative luciferase activities from reporters with or without the predicted dro-miR432 target sequence from *PGL3* in a dual-luciferase reporter assay using *Nicotiana benthamiana*. **(A)** Diagram of co-infiltrated sequences in the Agrobacterium tumefaciens–mediated infiltration assay. The reporter without the target sequence from *PGL3* (left) was co-infiltrated with each of two expression constructs (right): (1) the reporter without the target sequence and the *GUS* expression construct (Cont.–GUS) as a control, and (2) the reporter without the target sequence and the pri-miR432 expression construct (Cont.–miRNA). Similarly, the reporter with the target sequence from *PGL3* (left) was co-infiltrated with each of two expression constructs (right): (1) the reporter with the target sequence and the *GUS* expression construct (Target–GUS) as a control, and (2) the reporter with the target sequence and the pri-miR432 expression construct (Target–miRNA). **(B)** Relative F-Luc/R-Luc ratio after 48 h of incubation for the four co-infiltration pairs: Cont.–GUS, Cont.–miRNA, Target–GUS, and Target–miRNA. Different symbols in the same panels indicate the results of two independent experiments. The relative F-Luc/R-Luc ratios between two co-infiltrations were compared using Welch’s t-test. N.S., not significant; **, *p* < 0.01. The box boundaries represent the first (Q1) and third (Q3) quartiles, the horizontal line within each box indicates the median, and the whiskers extend to the most extreme data points within 1.5 × the interquartile range (IQR). **(C)** Relative F-Luc/R-Luc ratio after 72 h of incubation for the four co-infiltration pairs: Cont.–GUS, Cont.–miRNA, Target–GUS, and Target–miRNA. Different symbols in the same panels indicate the results of four independent experiments. The relative F-Luc/R-Luc ratios between two co-infiltrations were compared using Welch’s *t*-test. N.S., not significant; **, *p* < 0.01.

**Fig. S5.**
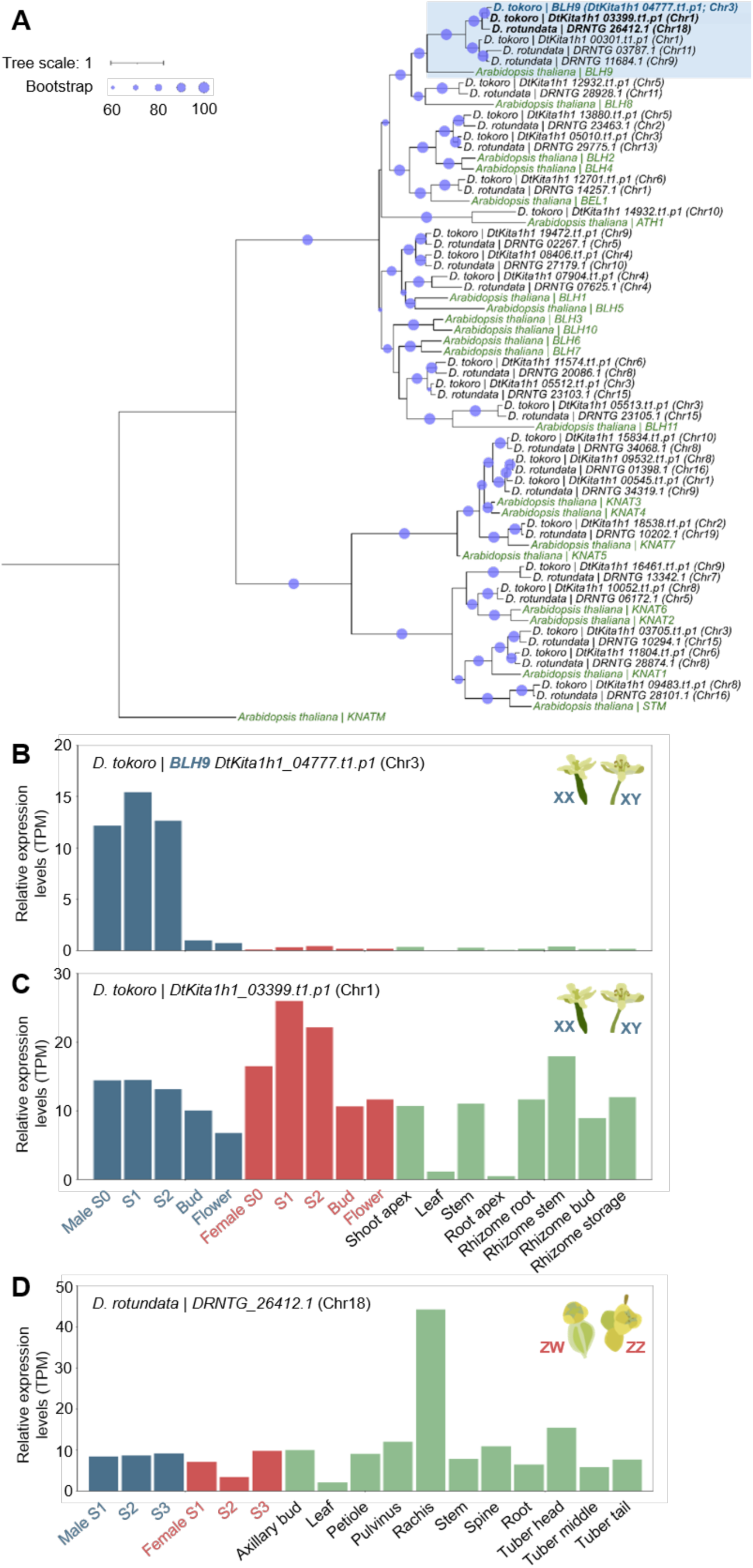
Relative expression levels of the putative sex determination gene *BLH9* in *Dioscorea tokoro* and its homologs. **(A)** Phylogenetic tree based on the protein sequences of BLH9 and TALE superfamily protein homologs from *D. tokoro* and *D. rotundata* (white Guinea yam), and TALE superfamily proteins from *Arabidopsis thaliana*. The circles in the tree nodes indicate bootstrap values. Tree scale indicates amino-acid substitutions per site. The relative expression levels of *BLH9* and putative homologs (shown in bold) are shown in panels (B)–(D). Each developmental stage and organ is represented by a single biological sample. **(B)** Relative expression levels of *BLH9* in male and female *D. tokoro* flowers at five stages of development and in non-reproductive organs. **(C)** Relative expression levels of a *BLH9* homolog from *D. tokoro*. **(D)** Relative expression levels of a *BLH9* homolog from *D. rotundata*.

**Fig. S6.**
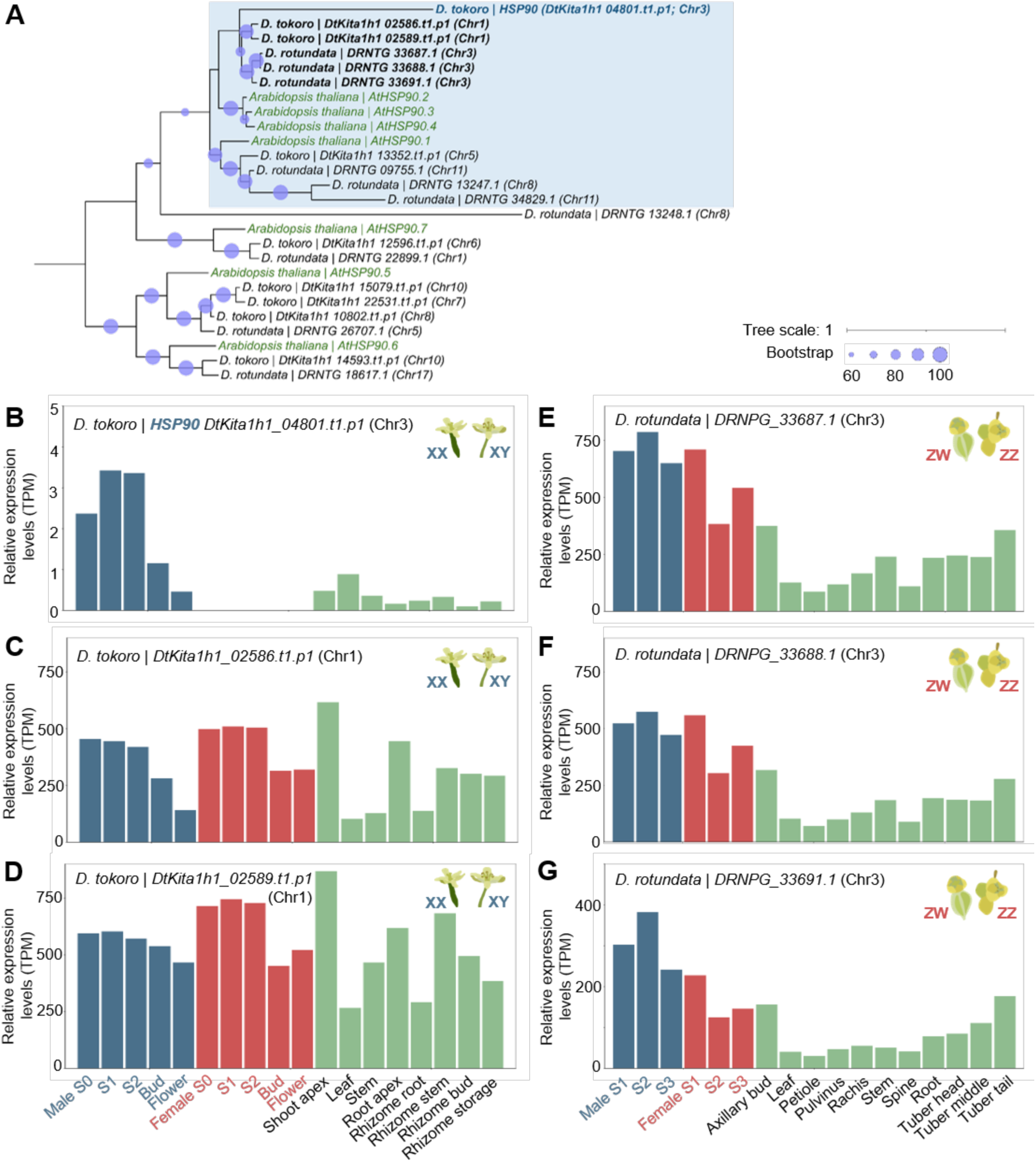
Relative expression levels of the putative sex determination gene *HSP90* in *Dioscorea tokoro* and its homologs. **(A)** Phylogenetic tree based on the cDNA sequences of *HSP90* homologs from *D. tokoro* and *D. rotundata* (white Guinea yam) and *AtHSP90* genes from *Arabidopsis thaliana*. The circles in the tree nodes indicate bootstrap values. Tree scale indicates nucleotide substitutions per site. The relative expression levels of *HSP90* and its putative homologs (shown in bold) are shown in panels (B)–(G). Each developmental stage and organ is represented by a single biological sample. **(B)** Relative expression levels of *HSP90* in *D. tokoro* male and female flowers at five stages of development and in non-reproductive organs. **(C,D)** Relative expression levels of putative *HSP90* homologs in *D. tokoro*. **(E–G)** Relative expression levels of putative *HSP90* homologs in *D. rotundata*.

**Fig. S7.**
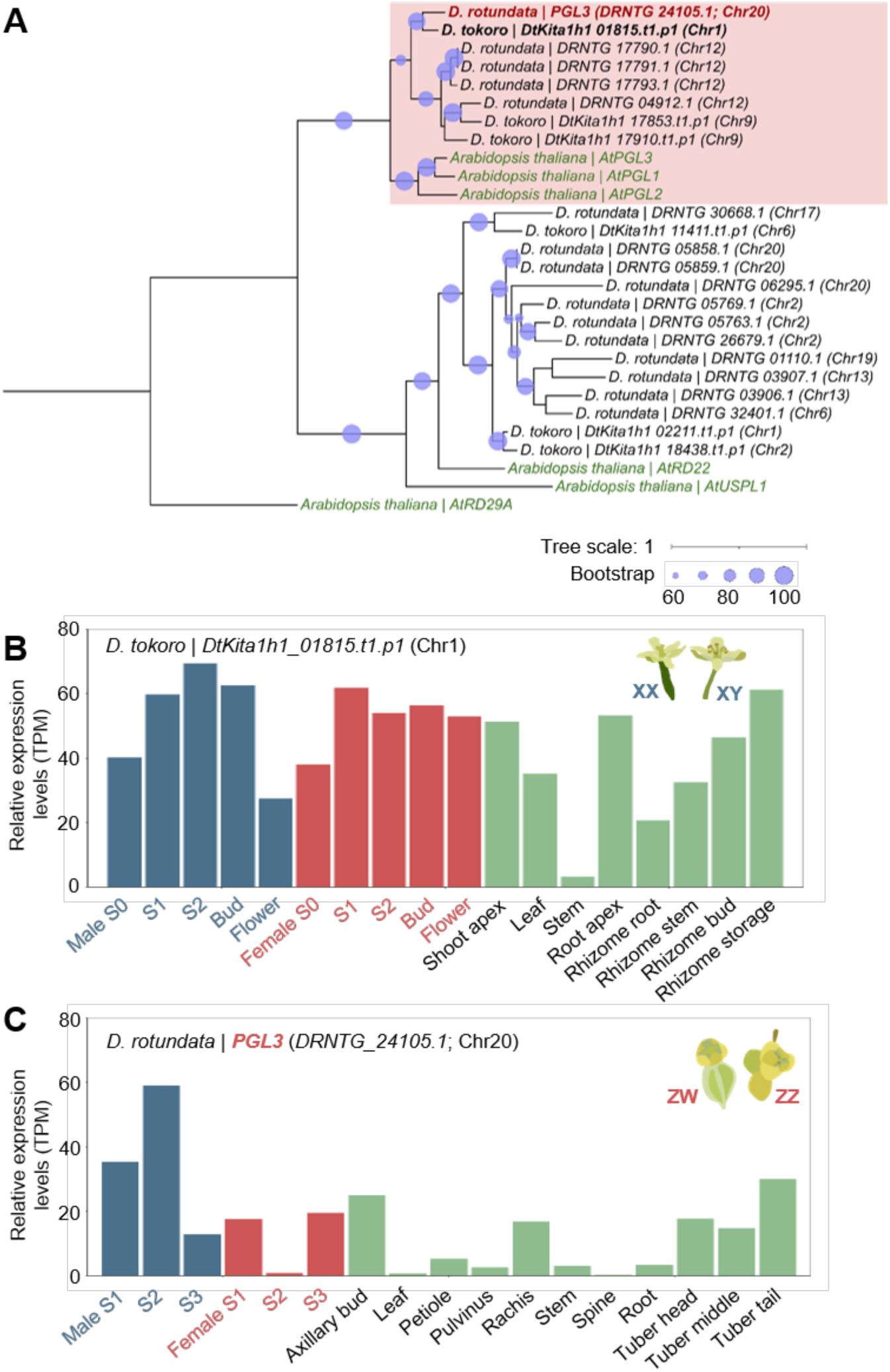
Relative expression levels of *PGL3*, the target gene of dro-miR432, in white Guinea yam, and its homologs. **(A)** Phylogenetic tree of the protein sequences of BURP homologs from *D. tokoro* and *D. rotundata*, and BURP domain–containing proteins from *Arabidopsis thaliana*. The circles in the tree nodes indicate bootstrap values. Tree scale indicates amino-acid substitutions per site. The relative expression levels of *PGL3* and putative homologs (shown in bold) are shown in panels (B)–(C). Each developmental stage and organ is represented by a single biological sample. **(B)** Relative expression levels of a *PGL3* homolog of *D. tokoro* in male and female flowers at five stages of development and in non-reproductive organs. **(C)** Relative expression levels of *PGL3* in *D. rotundat*

**Fig. S8.**
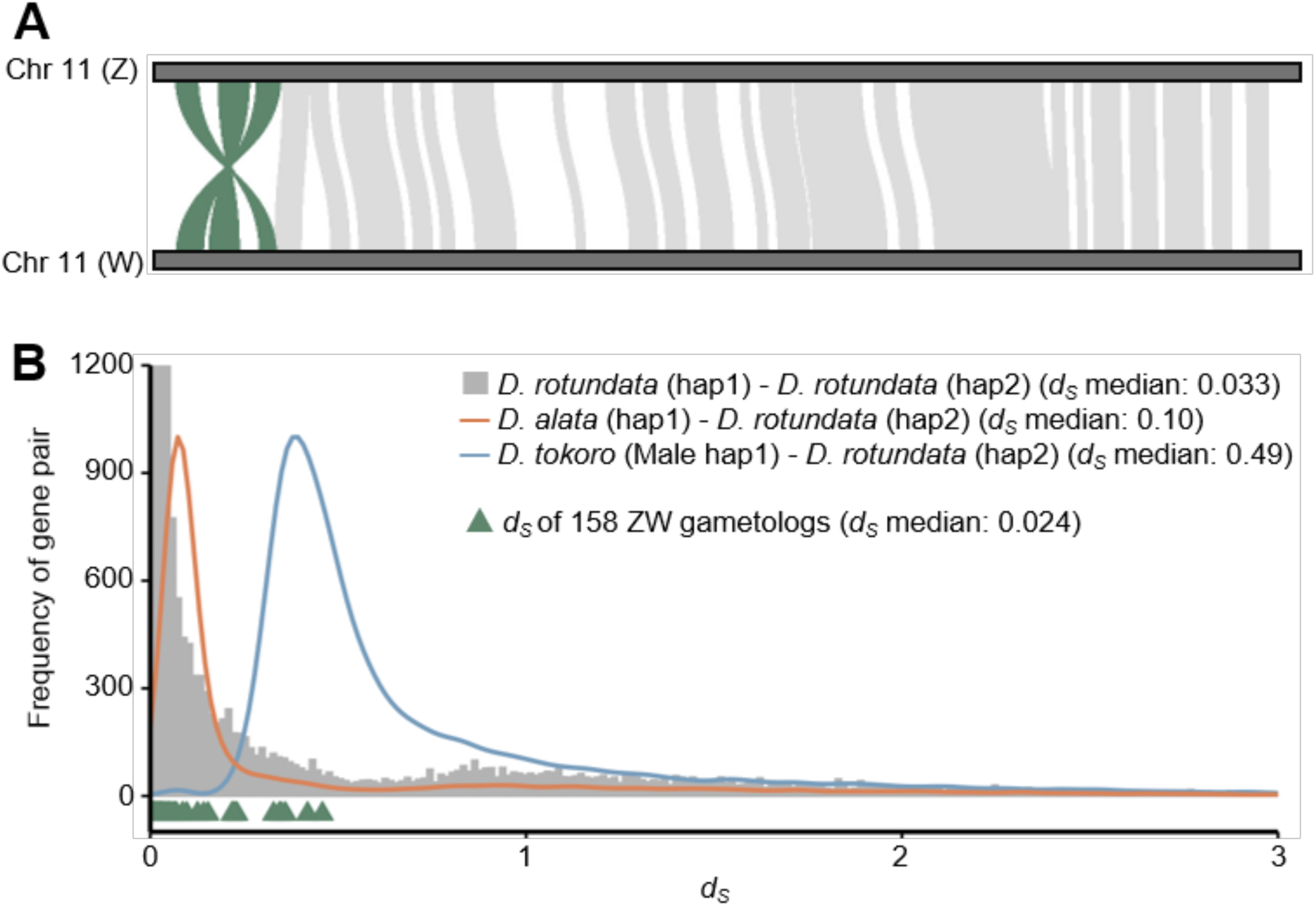
Distributions of intraspecific and interspecific synonymous site divergence (*dS*). **(A)** Chromosomal synteny analysis of the Z and W chromosomes also revealed inversions at the top of the sex chromosome. **(B)** Distribution of intraspecific *dS* values between all orthologous or paralogous gene pairs of the haplotype 1 and 2 assemblies of *D. rotundata* (bar graph in gray), and distributions of interspecific *dS* values of the *D. alata–D. rotundata* comparison (line graph in orange) and the *D. tokoro–D. rotundata* comparison (line graph in blue). Gene pairs were identified and their *dS* values calculated as described (*34–36*). All predicted genes (32,671 genes) with complete coding sequences from the *D. rotundata* haplotype 2 assembly were used as the reference, and 51,529 orthologous or paralogous gene pairs were identified by a comparison to the haplotype 1 assembly of *D. rotundata*, 36,014 gene pairs by a comparison to the *D. alata* genome, and 59,863 gene pairs by a comparison to the *D. tokoro* genome. A total of 158 genes in the inverted regions were defined as the Z–W gametologs. Inverted triangles indicate the *dS* values of gametologs.

**Fig. S9.**
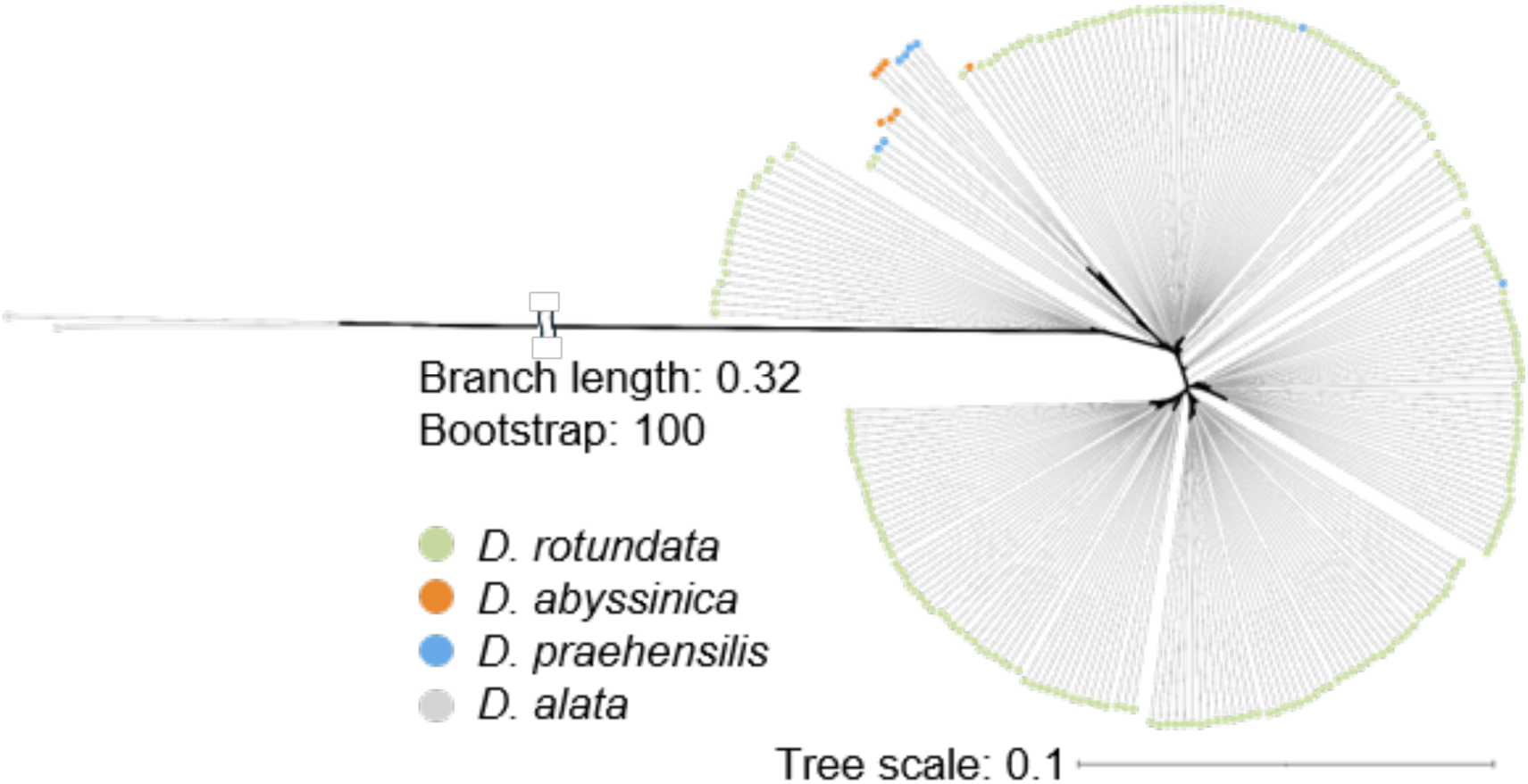
Phylogenetic tree of the W-specific region from white Guinea yam, *D. abyssinica*, and *D. praehensilis* using *D. alata* as an outgroup. Tree scale indicates nucleotide substitutions per site. A total of 219 white Guinea yam accessions, seven *D. abyssinica* accessions, and eight *D. praehensilis* accessions with sufficient coverage across the W-specific region were included in the phylogenetic analysis.

**Table S1.** Summary of the haplotype 1 and 2 genome assemblies.

| Assembly type |  | Haplotype-resolved contigs |  |
| --- | --- | --- | --- |
|  |  | Haplotype 1 | Haplotype 2 |
| *De novo* assembly | Total number of contigs | 1,209 | 280 |
|  | Total number of telomere-to-telomere contigs | 8 | 9 |
|  | Total number of base pairs (bp) | 630,575,853 | 570,381,407 |
|  | Average contig size (bp) | 521,568.1 | 2,037,076.5 |
|  | Longest contig (bp) | 40,755,161 | 37,062,405 |
|  | Shortest contig (bp) | 9,949 | 13,145 |
|  | N50 value (bp) | 17,927,676 | 15,137,522 |
| Genome evaluation<br>(BUSCO:<br>embryophyta_odb10) | Total BUSCO groups searched | 1,614 | 1,614 |
|  | Complete | 98.3% | 98.5% |
|  | Complete and single copy | 96.1% | 96.5% |
|  | Complete and duplicated | 2.2% | 2.0% |
|  | Fragmented | 0.6% | 0.6% |
|  | Missing | 1.1% | 0.9% |
| Gene annotation | Total number of lifted genes | 32,881 | 32,908 |
|  | (Total transcript variants) | (63,386) | (63,487) |

**Table S2.**
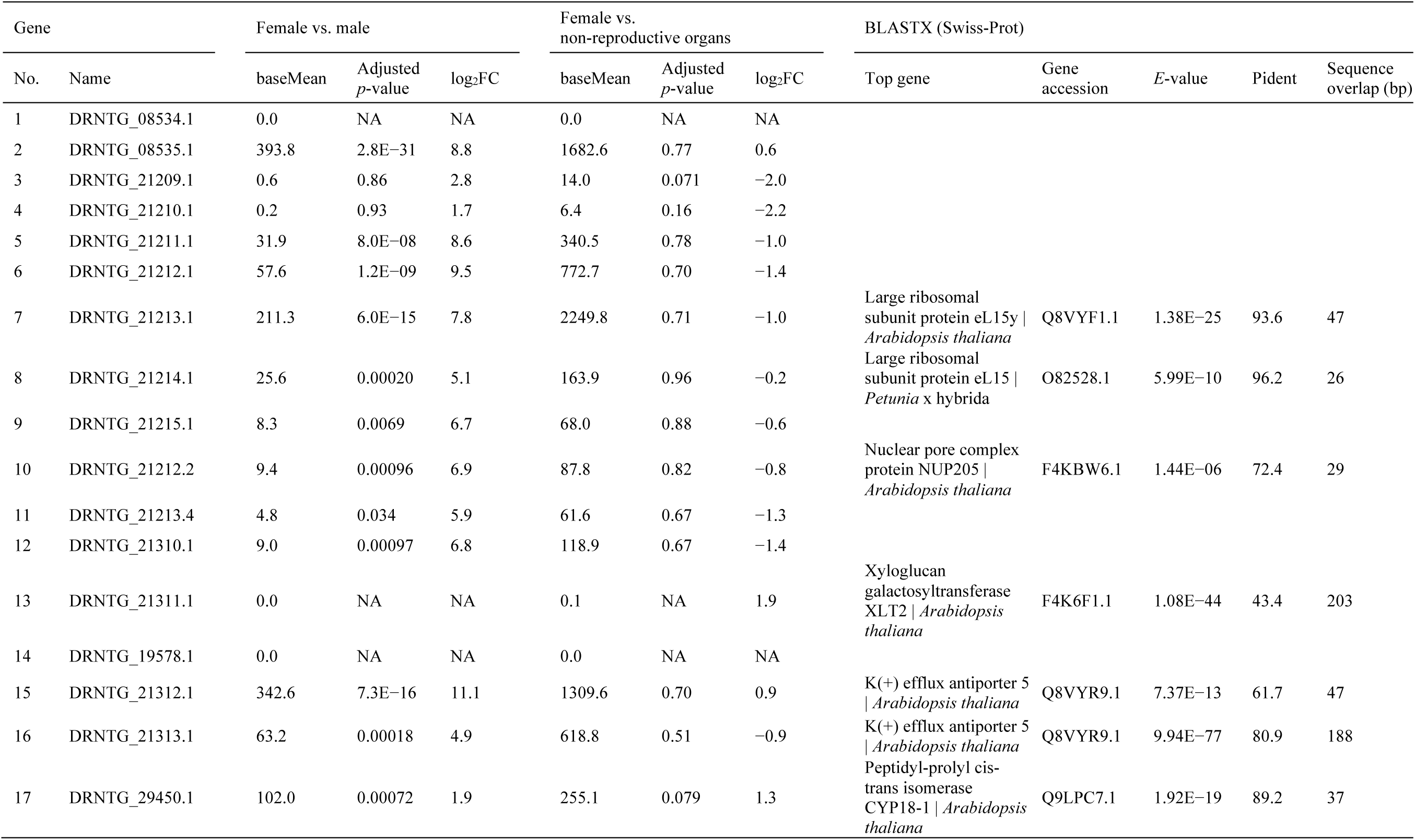
List of genes located in the W-specific region.

**Table S3.**
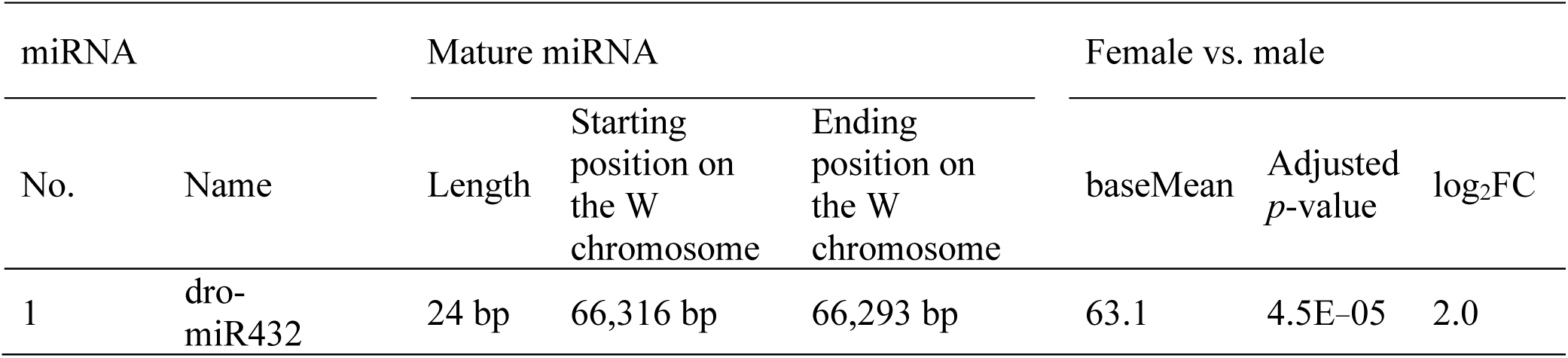
Information about the miRNA located in the W-specific region.

| miRNA |  | Mature miRNA |  |  | Female vs. male |  |  |
| --- | --- | --- | --- | --- | --- | --- | --- |
| No. | Name | Length | Starting position on the W chromosome | Ending position on the W chromosome | baseMean | Adjusted *p*-value | log<sub>2</sub>FC |
| 1 | dro-miR432 | 24 bp | 66,316 bp | 66,293 bp | 63.1 | 4.5E-05 | 2.0 |

**Table S4.** Sequence similarity search against the *D. rotundata* genome using genes located in the W-specific region.

| W-specific genes |  | BLASTP (*D. rotundata*) top hit excluding query transcripts |  |  |  |
| --- | --- | --- | --- | --- | --- |
| No. | Gene name | Gene name | Percent identity | Sequence overlap (aa) | Chromosome |
| 1 | DRNTG_08534.1 | DRNTG_05652.1 | 69.5 | 95 | chrom_5 |
| 2 | DRNTG_08535.1 | DRNTG_03226.1 | 48.7 | 117 | chrom_14 |
| 3 | DRNTG_21209.1 | DRNTG_21823.2 | 89.7 | 39 | chrom_18 |
| 4 | DRNTG_21210.1 | DRNTG_03642.1 | 80.4 | 56 | chrom_5 |
| 5 | DRNTG_21211.1 | – |  |  |  |
| 6 | DRNTG_21212.1 | – |  |  |  |
| 7 | DRNTG_21213.1 | DRNTG_05583.1 | 100 | 47 | chrom_16 |
| 8 | DRNTG_21214.1 | DRNTG_05583.1 | 100 | 26 | chrom_16 |
| 9 | DRNTG_21215.1 | – |  |  |  |
| 10 | DRNTG_21212.2 | – |  |  |  |
| 11 | DRNTG_21213.4 | – |  |  |  |
| 12 | DRNTG_21310.1 | – |  |  |  |
| 13 | DRNTG_21311.1 | DRNTG_06319.1 | 70.5 | 207 | chrom_13 |
| 14 | DRNTG_19578.1 | – |  |  |  |
| 15 | DRNTG_21312.1 | DRNTG_08535.3 | 100 | 89 | chrom_11 |
| 16 | DRNTG_21313.1 | – |  |  |  |
| 17 | DRNTG_29450.1 | DRNTG_01337.1 | 51.4 | 35 | chrom_13 |

**Table S5.** BLASTP analysis of the protein encoded by the target gene *PGL3*.

| Gene | BLASTP (Swiss-Prot) |  |  |  |  |
| --- | --- | --- | --- | --- | --- |
|  | Top hit gene | Gene accession | E-value | Pident | Query coverage (%) |
| *PGL3* | Polygalacturonase 1 beta-like protein 3 \| *Arabidopsis thaliana* | P92990.2 | 0 | 54.1 | 82 |
|  | Polygalacturonase-1 non-catalytic subunit beta \| *Solanum lycopersicum* | Q40161.1 | 0 | 53.3 | 80 |
|  | BURP domain-containing protein 12 \| *Oryza sativa* | Q6ZA27.1 | 0 | 54.7 | 80 |
|  | Polygalacturonase 1 beta-like protein 1 \| *Arabidopsis thaliana* | P92982.2 | 0 | 52.1 | 79 |
|  | Polygalacturonase non-catalytic subunit AroGP3 \| *Solanum lycopersicum* | P93218.1 | 0 | 52.1 | 79 |
|  | Polygalacturonase non-catalytic subunit AroGP2 \| *Solanum lycopersicum* | P93217.1 | 0 | 51.5 | 79 |
|  | Polygalacturonase 1 beta-like protein 2 \| *Arabidopsis thaliana* | O80760.1 | 0 | 50.3 | 84 |
|  | BURP domain-containing protein 14 \| *Oryza sativa* | Q6K2M1.2 | 0 | 49.0 | 79 |
|  | BURP domain-containing protein 16 \| *Oryza sativa* | Q7XES5.1 | 6.0E-80 | 58.0 | 32 |
|  | BURP domain-containing protein 5 \| *Oryza sativa* | Q0JEP3.2 | 6.0E-26 | 31.1 | 27 |

**Table S6.** Summary of PacBio HiFi long-read sequences and Omni-C sequences.

| Feature | PacBio<br>HiFi reads | Omni-C reads<br>(150 bp PE) |
| --- | --- | --- |
| Number of reads | 2,104,184 | 878,310,924 |
| Total length (bp) | 24,694,570,726 | 131,746,638,600 |
| Genome coverage | 43.3x | 231x |
| Average length (bp) | 11,736 | 150 |
| Longest length (bp) | 54,063 | — |
| N50 value (bp) | 12,173 | — |
| Accession number | SRR39875437 | SRR39875438 |

**Table S7.**
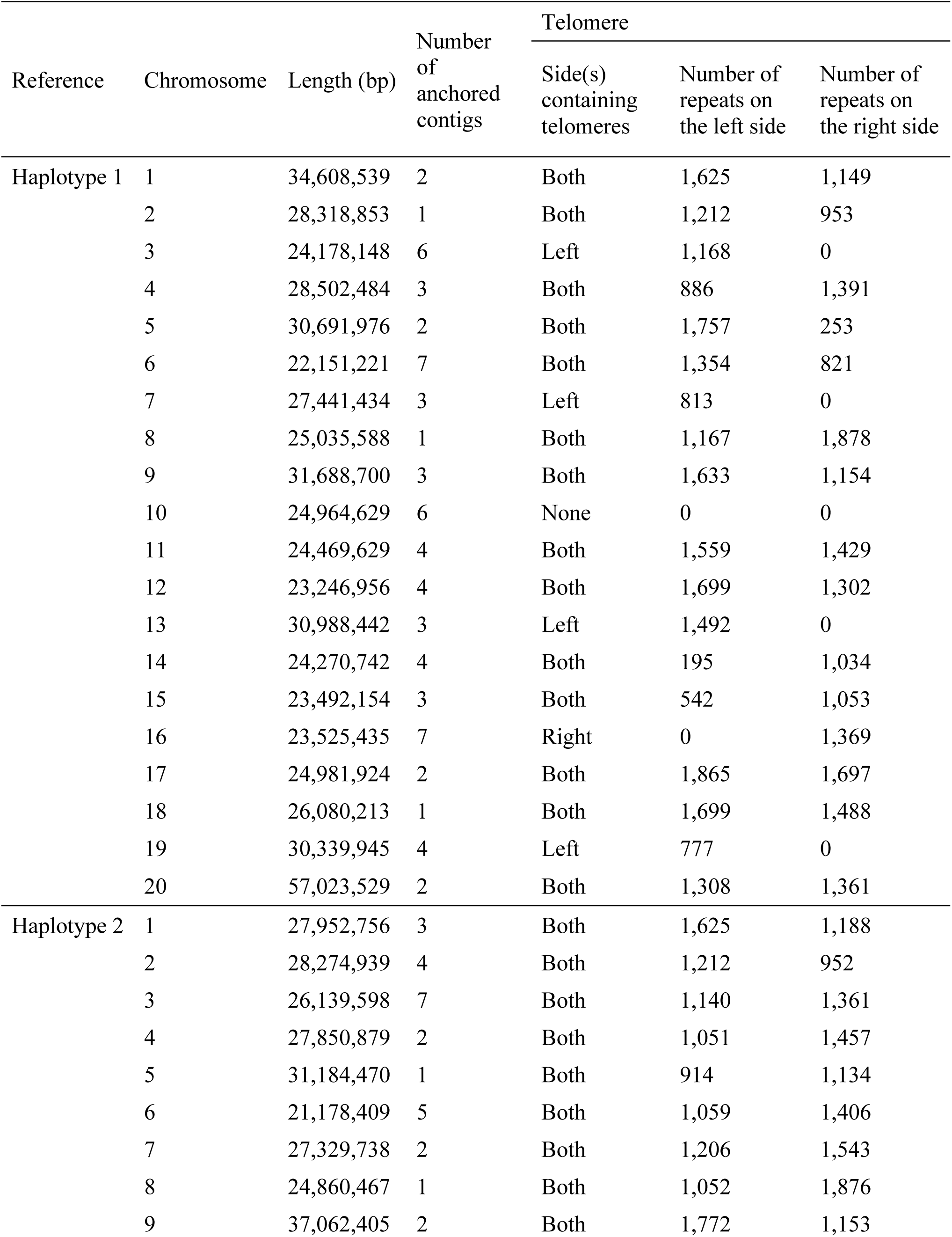
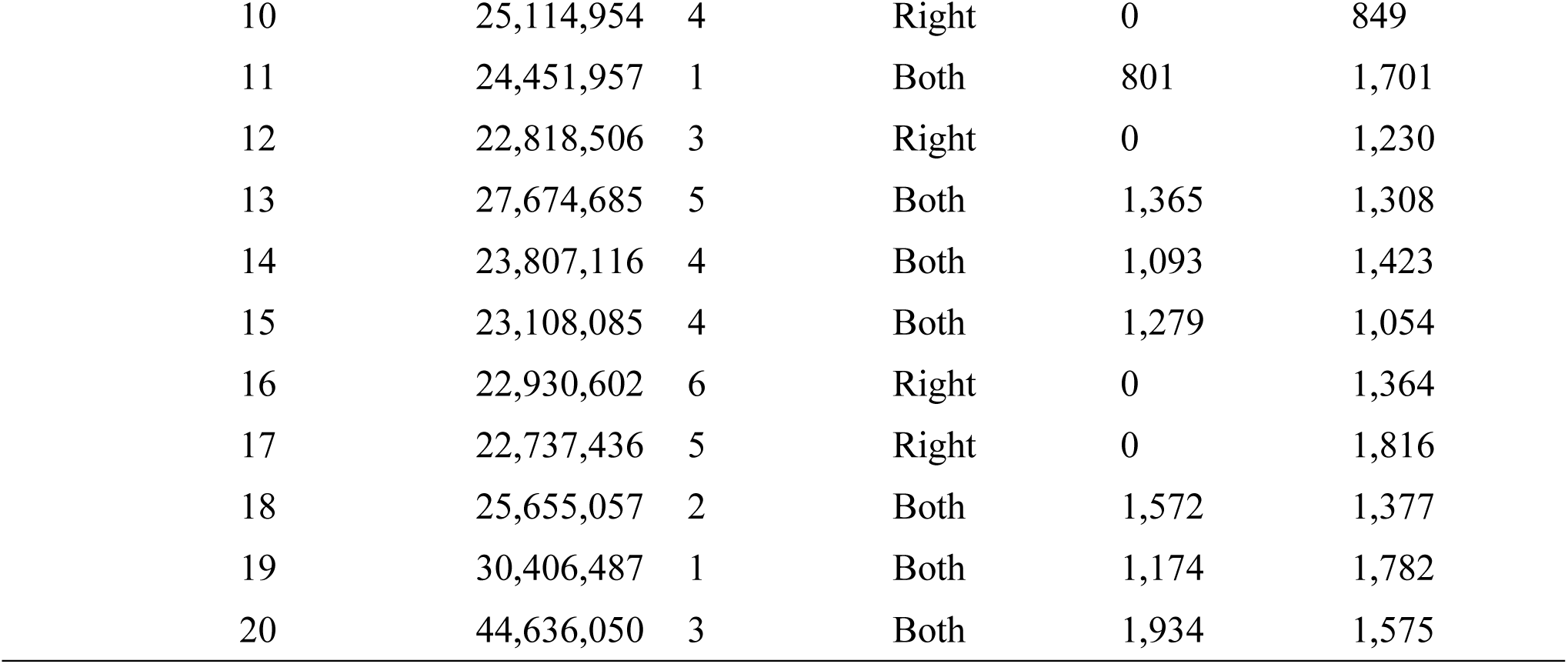
Number of anchored contigs and telomere repeats.

| Reference | Chromosome | Length (bp) | Number of anchored contigs | Telomere |  |  |
| --- | --- | --- | --- | --- | --- | --- |
|  |  |  |  | Side(s) containing telomeres | Number of repeats on the left side | Number of repeats on the right side |
| Haplotype 1 | 1 | 34,608,539 | 2 | Both | 1,625 | 1,149 |
|  | 2 | 28,318,853 | 1 | Both | 1,212 | 953 |
|  | 3 | 24,178,148 | 6 | Left | 1,168 | 0 |
|  | 4 | 28,502,484 | 3 | Both | 886 | 1,391 |
|  | 5 | 30,691,976 | 2 | Both | 1,757 | 253 |
|  | 6 | 22,151,221 | 7 | Both | 1,354 | 821 |
|  | 7 | 27,441,434 | 3 | Left | 813 | 0 |
|  | 8 | 25,035,588 | 1 | Both | 1,167 | 1,878 |
|  | 9 | 31,688,700 | 3 | Both | 1,633 | 1,154 |
|  | 10 | 24,964,629 | 6 | None | 0 | 0 |
|  | 11 | 24,469,629 | 4 | Both | 1,559 | 1,429 |
|  | 12 | 23,246,956 | 4 | Both | 1,699 | 1,302 |
|  | 13 | 30,988,442 | 3 | Left | 1,492 | 0 |
|  | 14 | 24,270,742 | 4 | Both | 195 | 1,034 |
|  | 15 | 23,492,154 | 3 | Both | 542 | 1,053 |
|  | 16 | 23,525,435 | 7 | Right | 0 | 1,369 |
|  | 17 | 24,981,924 | 2 | Both | 1,865 | 1,697 |
|  | 18 | 26,080,213 | 1 | Both | 1,699 | 1,488 |
|  | 19 | 30,339,945 | 4 | Left | 777 | 0 |
|  | 20 | 57,023,529 | 2 | Both | 1,308 | 1,361 |
| Haplotype 2 | 1 | 27,952,756 | 3 | Both | 1,625 | 1,188 |
|  | 2 | 28,274,939 | 4 | Both | 1,212 | 952 |
|  | 3 | 26,139,598 | 7 | Both | 1,140 | 1,361 |
|  | 4 | 27,850,879 | 2 | Both | 1,051 | 1,457 |
|  | 5 | 31,184,470 | 1 | Both | 914 | 1,134 |
|  | 6 | 21,178,409 | 5 | Both | 1,059 | 1,406 |
|  | 7 | 27,329,738 | 2 | Both | 1,206 | 1,543 |
|  | 8 | 24,860,467 | 1 | Both | 1,052 | 1,876 |
|  | 9 | 37,062,405 | 2 | Both | 1,772 | 1,153 |

|  |  |  |  |  |  |
| --- | --- | --- | --- | --- | --- |
| 10 | 25,114,954 | 4 | Right | 0 | 849 |
| 11 | 24,451,957 | 1 | Both | 801 | 1,701 |
| 12 | 22,818,506 | 3 | Right | 0 | 1,230 |
| 13 | 27,674,685 | 5 | Both | 1,365 | 1,308 |
| 14 | 23,807,116 | 4 | Both | 1,093 | 1,423 |
| 15 | 23,108,085 | 4 | Both | 1,279 | 1,054 |
| 16 | 22,930,602 | 6 | Right | 0 | 1,364 |
| 17 | 22,737,436 | 5 | Right | 0 | 1,816 |
| 18 | 25,655,057 | 2 | Both | 1,572 | 1,377 |
| 19 | 30,406,487 | 1 | Both | 1,174 | 1,782 |
| 20 | 44,636,050 | 3 | Both | 1,934 | 1,575 |

**Table S8.**
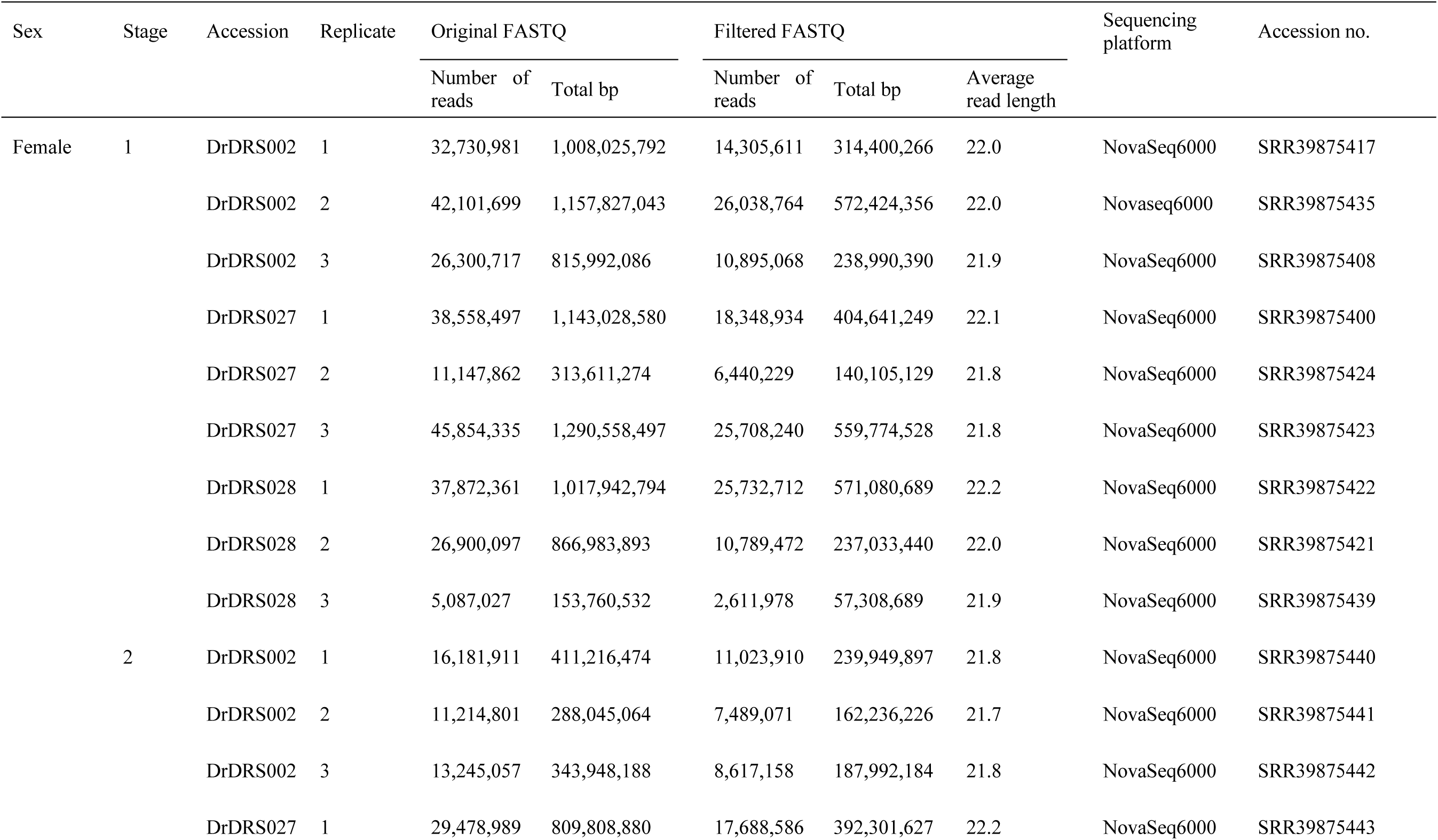
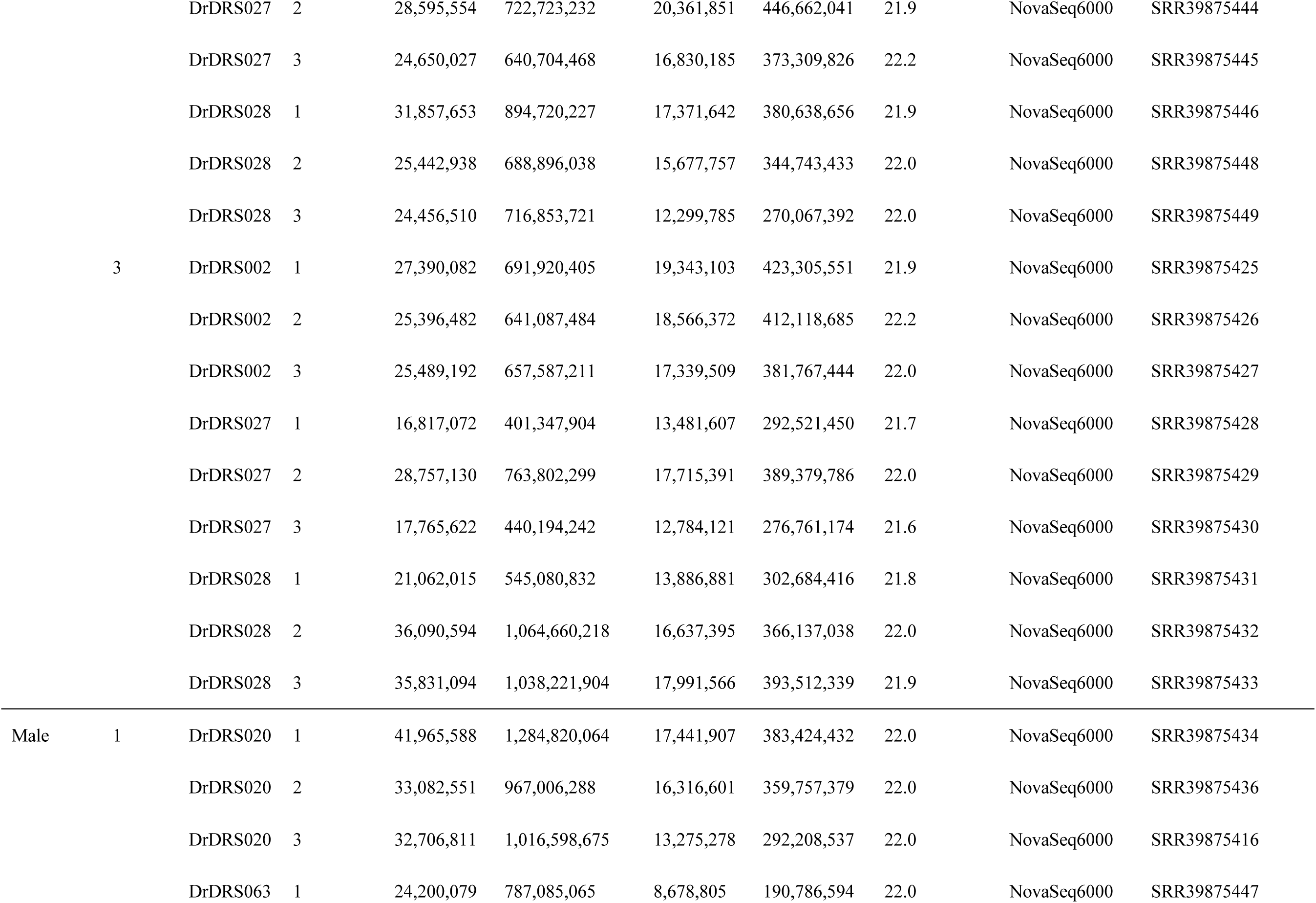
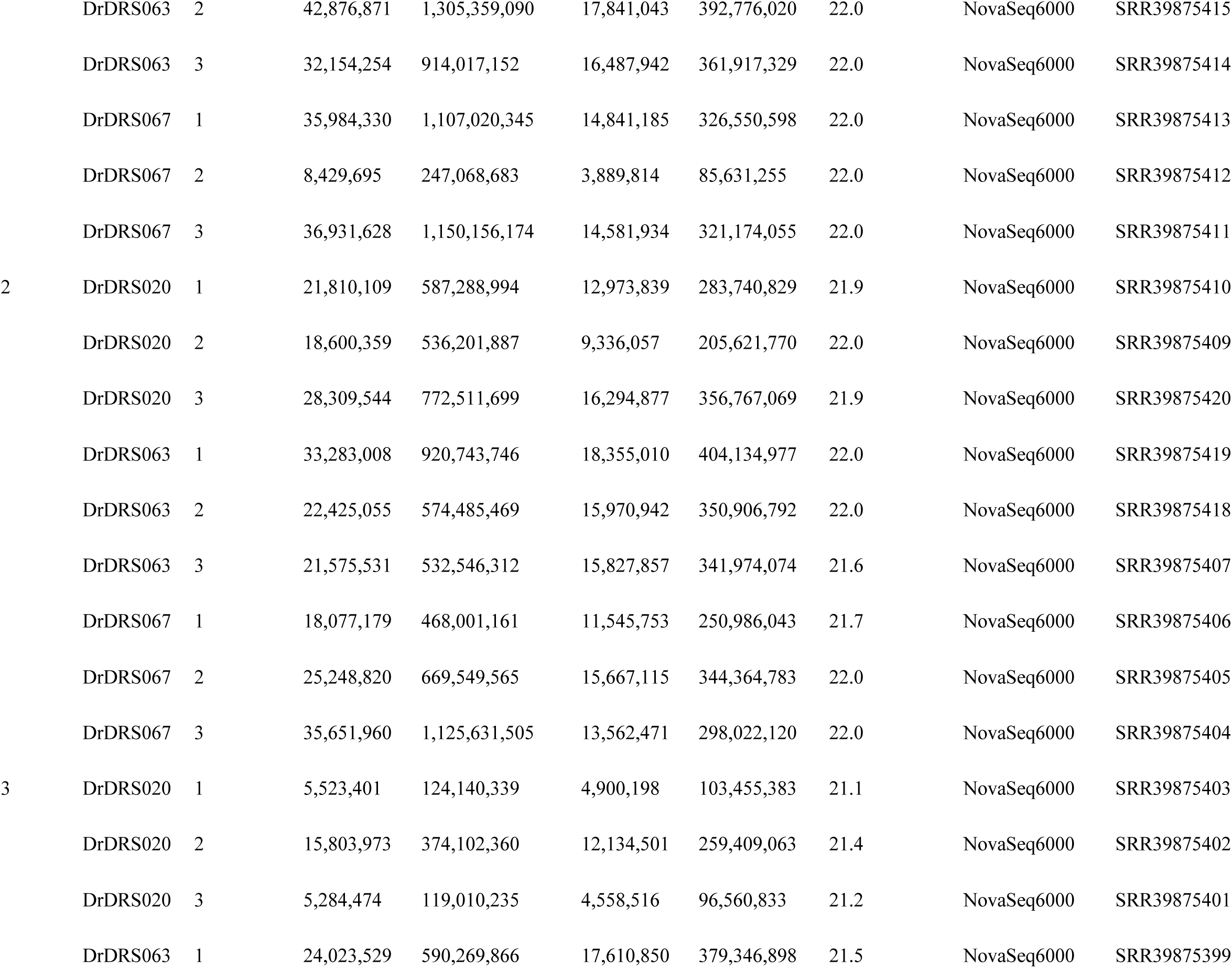
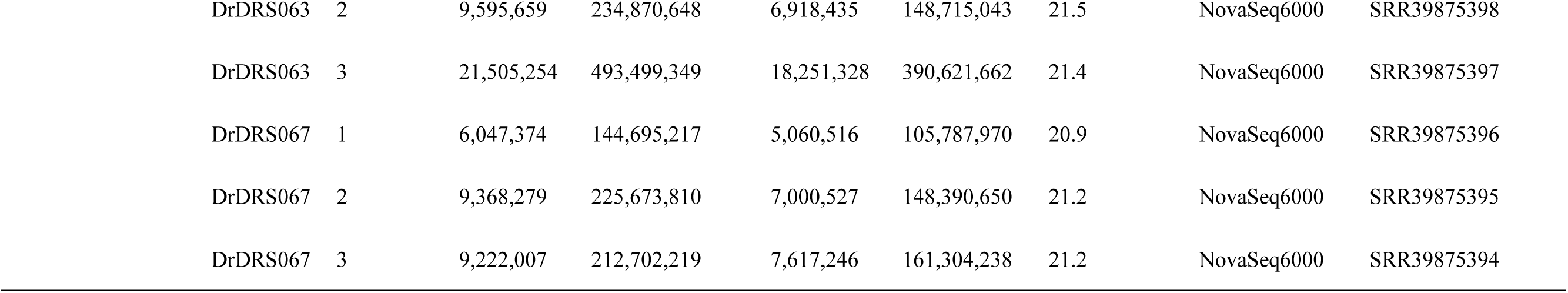
Small RNA-seq data generated from libraries constructed using female or male *Dioscorea rotundata* flowers at three stages of development.

**Table S9.**
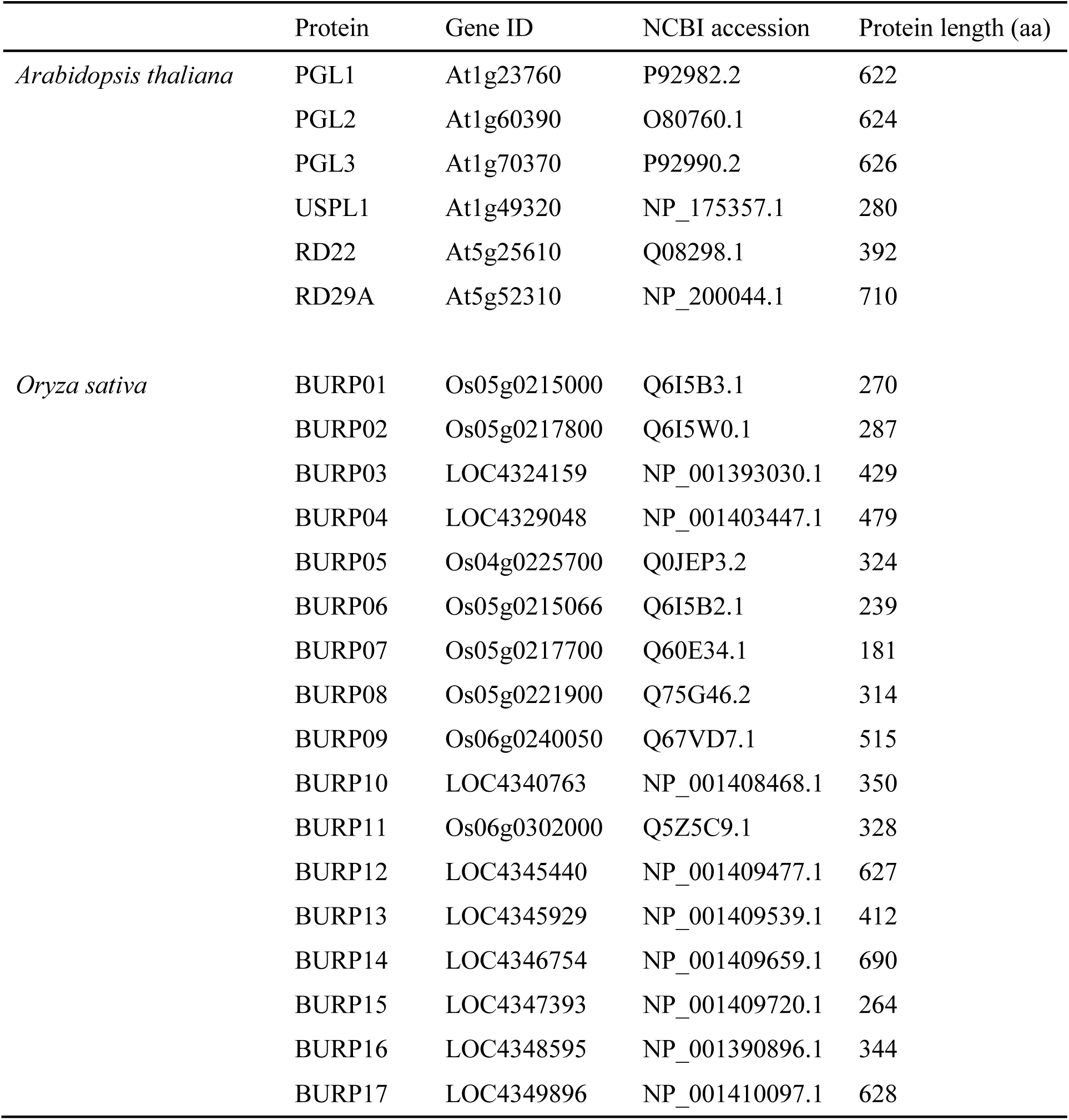
BURP domain proteins in *Arabidopsis thaliana* and *Oryza sativa* obtained from NCBI.

**Table S10.**
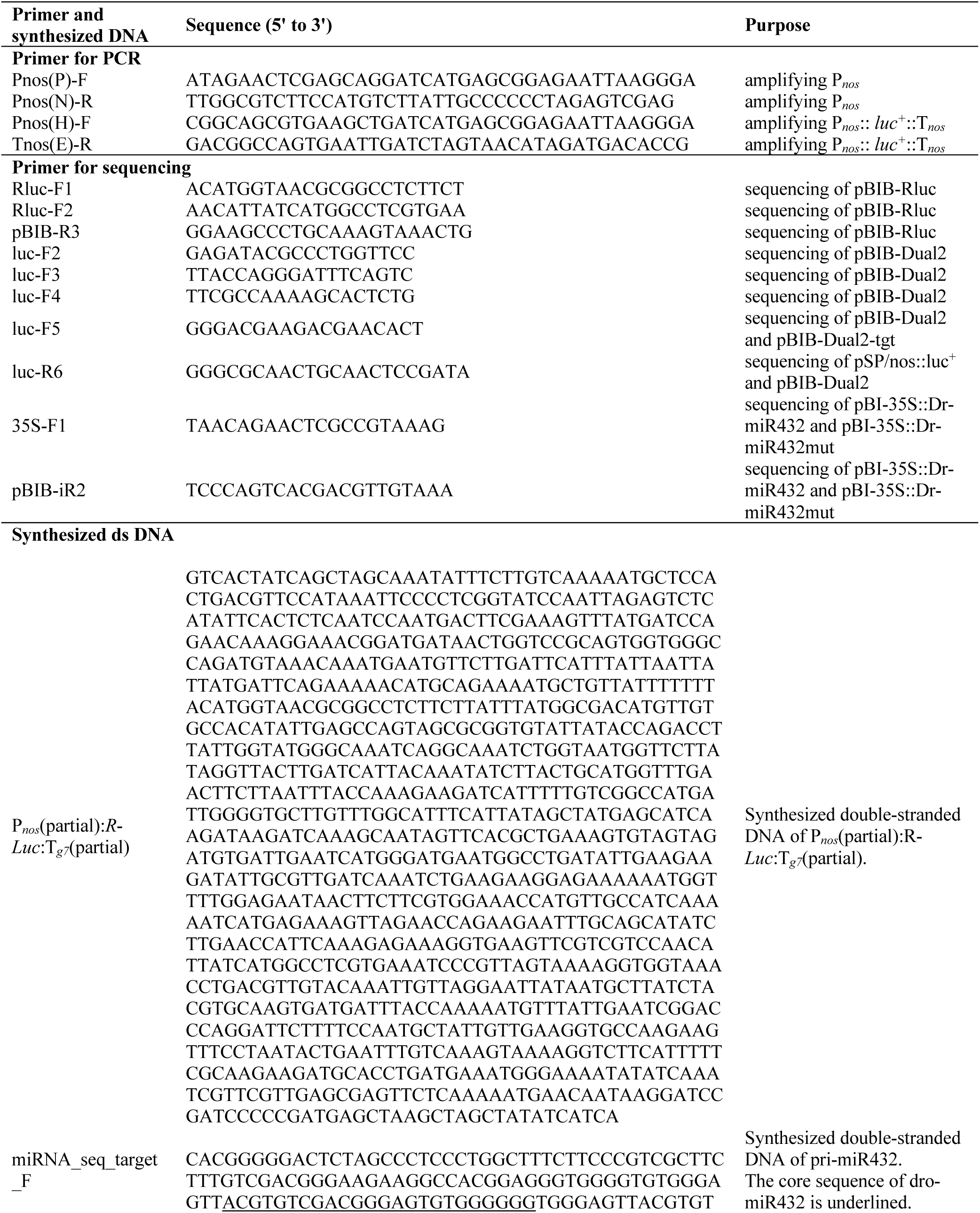
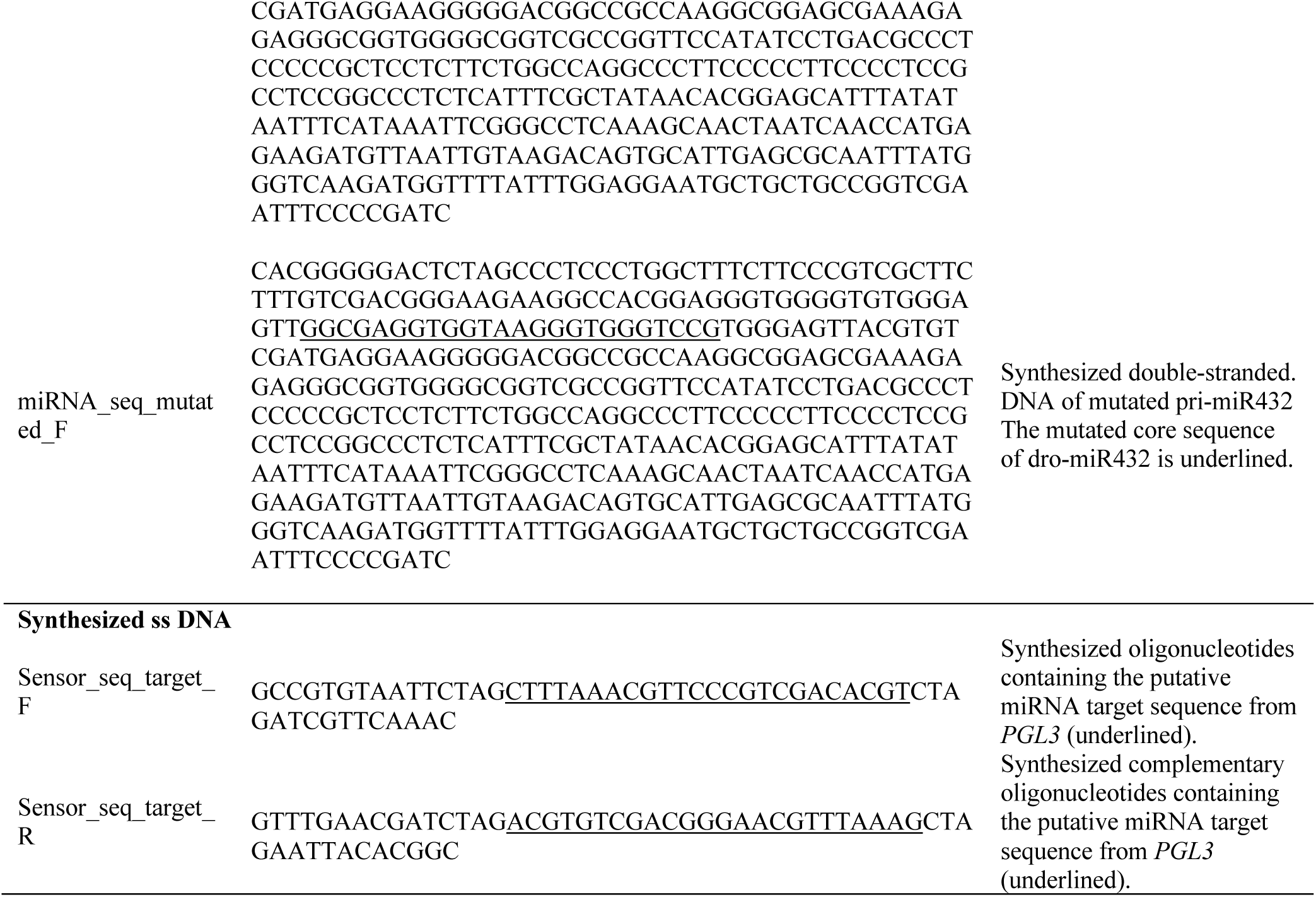
Primers and synthesized DNAs used in the dual-luciferase reporter assay in *Nicotiana benthamiana* leaves.

**Data S1. (separate file)**

Raw bioluminescence data for F-Luc and R-Luc in the dual-luciferase reporter assay in *Nicotiana benthamiana* leaves using three expression constructs: Target-GUS, Target-miRNA, and Target-Mutated (Fig. 3, fig. S3).

**Data S2. (separate file)**

Raw bioluminescence data for F-Luc and R-Luc in the dual-luciferase reporter assay in *Nicotiana benthamiana* leaves using four expression constructs: Cont.-GUS, Cont.-miRNA, Target-GUS and Target-miRNA (fig. S4).

**Data S3. (separate file)**

Illumina short reads from 332 *Dioscorea rotundata* accessions.

**Data S4. (separate file)**

Illumina short reads from 29 *Dioscorea abyssinica* accessions, 39 *Dioscorea praehensilis*accessions and two *Dioscorea alata* accessions.

